# Metabolites produced by gut bacteria under anoxic conditions drive the suppression of Acinetobacter baumannii

**DOI:** 10.1101/2025.09.29.676038

**Authors:** Katrin Anja Winter, Lisa Osbelt, Till Robin Lesker, Marie Wende, Éva d. H. Almási, Lea Eisenhard, Leonard Knegendorf, Birte Abt, Jörg Overmann, Michael Hogardt, Volkhard A. J. Kempf, Dirk Schlüter, Meina Neumann-Schaal, Till Strowig

## Abstract

Multidrug-resistant (MDR) bacteria pose a significant global health threat. Among these, *Acinetobacter baumannii,* particularly carbapenem-resistant *A. baumannii*, is a leading cause of healthcare-associated infections. Emerging evidence links gut colonization to systemic infections, highlighting opportunities for new control strategies. We demonstrate that *A. baumannii* can survive under anaerobic conditions (≤1%) and shows limited growth under low oxygen conditions (≥1%), underscoring its adaptation to niches in the intestine. However, specific carbohydrates, including maltose, provide the gut bacteria *Klebsiella oxytoca* with a competitive advantage under anoxic conditions, enabling it to actively suppress *A. baumannii* through its metabolism. Notably, maltose induces this suppressive capacity in other commensals and complex gut microbiomes as well. Force-feeding experiments in *Galleria mellonella* larvae corroborate that *K. oxytoca* in combination with maltose significantly reduces *A. baumannii* recovery in gut environments. These findings suggest that targeted carbohydrate supplementation could enhance probiotic strategies, creating an environment unfavorable to *A. baumannii*.

## Introduction

Multidrug-resistant (MDR) bacteria pose a significant global health threat, particularly due to their prevalence in clinical environments and ability to evade even last-line antibiotics ^1,2^. The World Health Organization (WHO) has identified the ESKAPE pathogens-*Enterococcus faecium, Staphylococcus aureus, Klebsiella pneumoniae, Acinetobacter baumannii, Pseudomonas aeruginosa*, and *Enterobacter* species-as the most critical MDR organisms, emphasizing their status as a top priority for monitoring and control efforts ^3–5^. Among these, *Acinetobacter baumannii*, especially carbapenem-resistant strains (CRAB), has emerged as a leading cause of healthcare-associated infections, resulting in pneumonia, bloodstream infections, and wound infections, particularly in immunocompromised patients ^6,7^. The increasing resistance of *A. baumannii* to carbapenems, one of the most effective classes of antibiotics^8,9^, is particularly concerning in healthcare settings, where the pathogen thrives and persists in intensive care units (ICUs) and other high-risk areas, causing nosocomial outbreaks that are difficult to control ^10–13^.

The human microbiome, particularly the intestinal microbiome, plays a vital role in health and disease, influencing various physiological functions such as metabolism, immune responses, and protection against pathogens ^14–16^. It comprises diverse, complex ecosystems where niche competition and bacterial interactions are crucial for maintaining homeostasis ^17–20^. Due to its high microbial density, the gut microbiome features a multiplicity of bacterial interactions, facilitating genetic exchanges, metabolic cooperation, and competition for resources, along with various direct or indirect agonistic and antagonistic interactions ^21–29^. These interactions contribute to critical microbiome functions like colonization resistance, which limits the ability of invading pathogens to establish themselves in this niche^30–32^. Alterations in microbiome composition, such as those induced by antibiotic use, can disrupt these complex protective mechanisms, thereby increasing gut colonization by resistant pathogens and, consequently, susceptibility to infections ^30,31,33–35^.

Members of the *Klebsiella oxytoca* species complex (KoSC) have traditionally been recognized as opportunistic pathogens ^36–39^. Recently, they have been acknowledged as key members of the microbiota, protecting the host from pathogens such as *Klebsiella pneumoniae* ^40,41^ and *Salmonella enterica* ^42^ through nutrient competition^43^, toxin production, and bacteriocin secretion. Additionally, niche exclusion by *K. oxytoca* can limit colonization by *K. pneumoniae*, a well-known cause of pneumonia and bloodstream infections ^40^, and its presence has been associated with reduced risks of bacteremia for cancer patients^44^. Based on these and other studies demonstrating anti-inflammatory effects of KoSC ^43^, efforts are underway to explore these bacteria as a next-generation probiotic. Notably, while the interplay between *K. pneumoniae* and *A. baumannii,* which frequently co-colonize the respiratory and gastrointestinal tracts ^45–53^, have been studied within polymicrobial infections ^53–57^, the crosstalk between *K. oxytoca* and *A. baumannii* has not yet been investigated.

In this study, we aimed to explore the interaction between *K. oxytoca* and CRAB strains, focusing on the impact of environmental conditions, such as oxygen availability and the presence of sugars, on their coculture dynamics. We identified that under oxygen-deprived conditions, *K. oxytoca* exerts strong inhibitory activity against *A. baumannii*. Specifically, under anoxic conditions, commonly found in the homeostatic intestine, diverse carbohydrates provide *K. oxytoca* with a strong advantage in suppressing *A. baumannii*. Transwell and spent media experiments suggested a contact-independent mechanism, and genetic loss-of-function experiments demonstrate that this effect is independent of *K. oxytoca*’s previously identified antimicrobial toxins. Notably, *in vivo* experiments in *Galleria mellonella* larvae confirmed that *K. oxytoca* in combination with maltose markedly reduces *A. baumannii* colonization. These results highlight the significance of metabolic interactions within the microbiome and suggest potential strategies for managing *A. baumannii* colonization in clinical settings.

## Results

### *K. oxytoca* suppresses *Acinetobacter* under anoxic conditions in the presence of specific carbon sources

Nutrient and, more broadly niche competition between pathogens and commensal bacteria or probiotic candidates are important in determining the outcome of microbial interactions. As representative models to evaluate *A. baumannii* and *K. oxytoca* interactions under controlled conditions, we utilized a CRAB strain (FR4326) isolated from human feces and a *K. oxytoca* strain with antagonistic properties against *K. pneumoniae* and other enterobacteria^40,42^. To compare the metabolic versatility of the two strains, the Biolog Phenotype MicroArray system using PM1 and PM2A plates was employed under aerobic, dye-free conditions to enable robust growth. The optical density at 600 nm (OD_600_) was continuously monitored for 24 hours, and only carbon sources (CS) with ODmax>0.2 were considered for subsequent analysis. Growth was analyzed by calculating the area under the curve (AUC) for each carbon source (**Fig. 1a**). The two strains demonstrated the capacity to utilize a distinct range of carbon sources (*A. baumannii* 22/190 CS, *K. oxytoca* 61/190 CS), with a subset of 16 CS shared between the strains (**Fig. 1b**). Notably, *K. oxytoca* showed more substantial growth on various carbohydrates, particularly simple sugars, while *A. baumannii* favored organic and amino acids, suggesting that these bacteria occupy different metabolic niches.

**Fig. 1:**
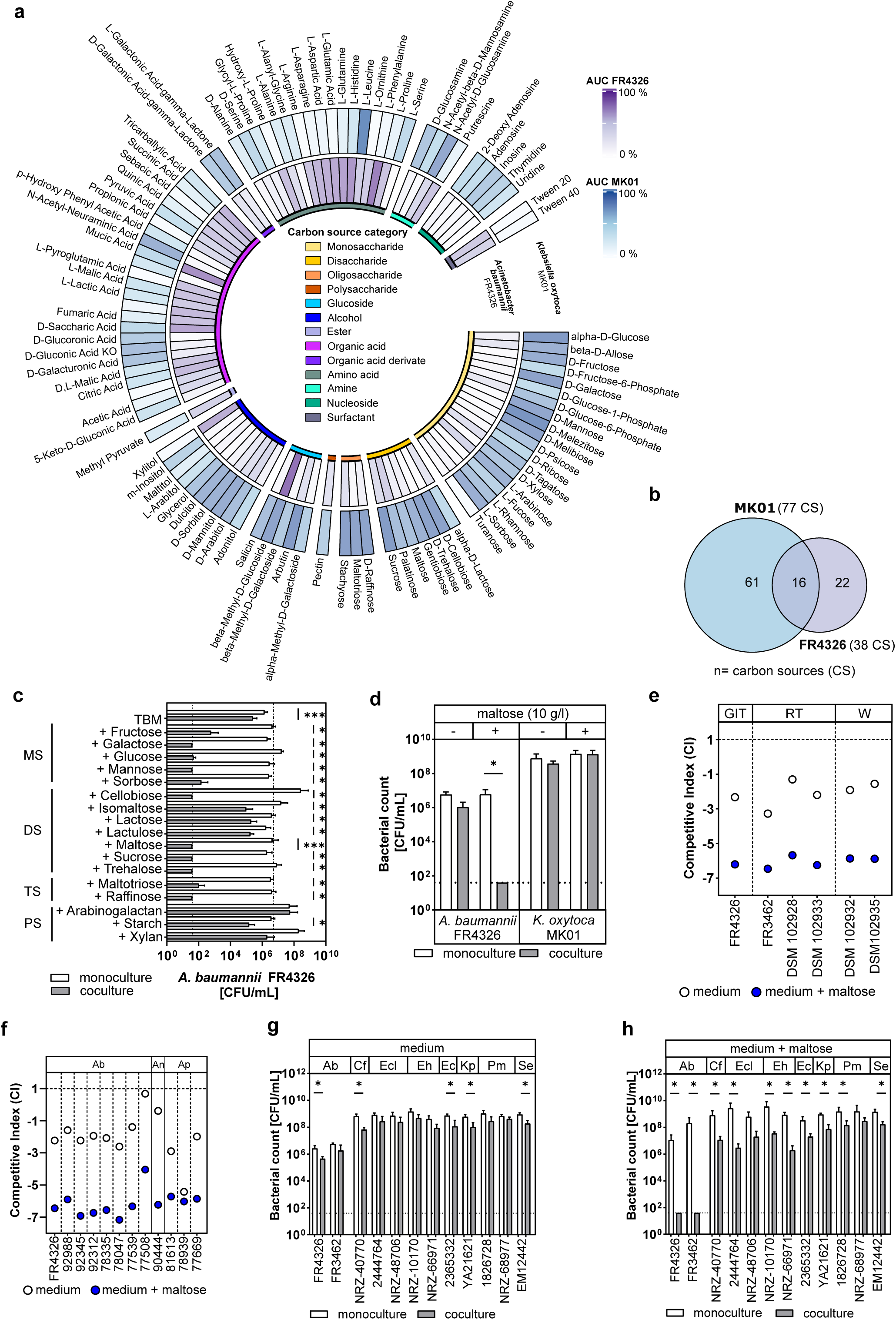
Carbon source-dependent growth profiles of *Klebsiella oxytoca* MK01 and *Acinetobacter baumannii* FR4326 in monoculture and coculture conditions. **a, b** *Klebsiella oxytoca* MK01 and *Acinetobacter baumannii* FR4326 were cultured in Biolog Phenotype MicroArray PM1 and PM2A plates. (**a**) The circular heatmap displays the area under the curve (AUC) of OD_600_ measurements for the indicated carbon sources. Only carbon sources that support growth (OD_600_ > 0.2) are shown for n= 3 biological replicates. (**b**) Number of unique and shared carbon sources with detected growth for *K. oxytoca* MK01 and A*. baumannii* FR4326. Growth was defined as OD_600_ > 0.2 after 24 hours of incubation. **c** *A. baumannii* FR4326 was cultured alone or in coculture with *K. oxytoca* MK01 in tryptone base medium (TBM) supplemented with indicated carbohydrates, including monosaccharides (MS), disaccharides (DS), trisaccharides (TS) and polysaccharides (PS) at a concentration of 10 g/L. The growth of *A. baumannii* was determined by selective plating. Data are presented as means ± standard deviation (SD) for n= 4 biological replicates. Assay controls (“TBM” and “+Maltose”) were conducted in n= 8 biological replicates. **d** *A. baumannii* FR4326 and *K. oxytoca* MK01 were cultured in mono- and cocultures in TBM ± 10 g/L maltose. The growth of *A. baumannii* and *K. oxytoca* was determined by selective plating. The mean ± SD for n= 4 biological replicates is displayed. **e**, f *K. oxytoca* strain MK01 was cocultured with the indicated *Acinetobacter* spp. strains in TBM ± 10 g/L maltose. (**f**) *A. baumannii* strains from the gastrointestinal tract (GIT), respiratory tract (RT), and skin wounds (W) (**g**) GIT isolates of *A. baumannii* (Ab), *A. nosocomialis* (An), and *A. pittii* (Ap). The growth of *Acinetobacter* strains and *K. oxytoca* was assessed by selective plating and the competitive index was calculated. Coculture experiments were conducted for n= 3 (**f**) and n= 4 (**g**) biological replicates. **g**, h *K. oxytoca* MK01 was cocultured with the indicated MDR enterobacteria in TBM, both without **(h)** and with 10 g/L maltose (**i**). The growth of MDR enterobacteria and *K. oxytoca* was assessed through selective plating. The mean ± SD for n= 4 biological replicates is displayed. *A. baumannii* (Ab) FR4326, FR3462; *C. freundii* (Cf) NRZ-40770; *E. cloacae* (Ecl) 2444764, NRZ-48706; *E. hormaechei* (Eh) NRZ-10170, NRZ-66971; *E. coli* (Ec) 2365332; *K. pneumoniae* (Kp) YA21621; *P. mirabilis* (Pm) 1826728, NRZ-68977; *S. enterica* (Se) SL1344. All coculture assays were incubated under oxygen-limited conditions using the AnaeroGen system. The detection limit is indicated (dotted line). Statistical significance was determined using Mann-Whitney U test (*p < 0.05).

Next, to study the interactions between *K. oxytoca* and *A. baumannii*, we established a simple coculture system under the following conditions: i) Cocultures were inoculated with a 10:1 ratio of *K. oxytoca* to *A. baumannii*, simulating a scenario of an established community with a higher abundance of *K. oxytoca* and a lower abundance of the invading pathogen; ii) Cocultures were cultivated in tryptone base medium (TBM), with and without various carbohydrates (10 g/L), to provide sufficient preferred nutrients for both bacteria; iii) Cocultures were incubated for 24 hours using an AnaeroGen system, which creates an anoxic environment similar to a homeostatic intestine, followed by selective plating to determine the numbers of viable bacteria for each species.

The coculture of *K. oxytoca* and *A. baumannii* in TBM did not adversely affect the survival of *A. baumannii* compared to the monocultures (Fig. 1c). In contrast, in TBM supplemented with various carbon sources such as galactose, glucose, mannose, maltose, and sucrose, the survival of *A. baumannii* was strongly reduced (>10^4^-fold reduction) only when *K. oxytoca* was present (**Fig. 1c**). Notably, other carbohydrates like lactose, lactulose, isomaltose, and starch did not lead to a comparable reduction in *A. baumannii,* despite similar growth of *K. oxytoca* in both inhibiting and neutral conditions observed with maltose supplementation. Maltose supplementation consistently reduced the levels of *A. baumannii* to the detection limit of the assay (**Fig. 1d; Supplementary Fig. 1a**). Therefore, we used this carbohydrate to test the robustness of the inhibitory effect extensively. Specifically, we varied the inoculation ratio and maltose concentrations. Remarkably, even at an inoculation ratio of *K. oxytoca* to *A. baumannii* of 1:1000, suppression of *A. baumannii* was observed (**Supplementary Fig. 1b, 1c**). Low maltose concentrations (1 g/L) already induced a significant reduction in *A. baumannii*, but higher concentrations (>5 g/L) were needed for complete suppression (**Supplementary Fig. 1d**).

To investigate whether the suppressive effect of *K. oxytoca* on *A. baumannii* was specific to the *A. baumannii* strain FR4326, the coculture assay was repeated with additional *Acinetobacte*r strains in TBM, both without and with maltose supplementation. First, five additional *A. baumannii* strains originating from the respiratory tract, skin wounds or lacerations were tested. A similar maltose-dependent suppressive effect on these *A. baumannii* strains was observed (**Fig. 1e; Supplementary Fig. 1e**). Next, we tested further *Acinetobacter* spp. isolated from the human intestine, including seven strains of *A. baumannii*, one strain of *A. nosocomialis,* and three strains of *A. pittii*. Strikingly, viable bacterial counts of all tested *Acinetobacter* spp. strains were reduced by up to 10^4^-fold in cocultures in a maltose-dependent manner (**Fig. 1f; Supplementary Figs 1f-1k**).

Finally, to investigate whether similar maltose-dependent reductions occur in the coculture of *K. oxytoca* with other (opportunistic) pathogens, assays were repeated with a panel of enterobacteria and enteropathogens. This panel included clinical isolates of *Citrobacter freundii*, *Enterobacter cloacae*, *Enterobacter hormaechei*, *Escherichia coli*, *Klebsiella pneumoniae*, *Proteus mirabilis*, and *Salmonella enterica,* all isolated from the human intestine. Coculture assays were conducted in TBM with and without maltose as the added carbon source. In TBM without maltose, the coculture with *K. oxytoca* resulted in no to modest reductions (<10^2^-fold) of the pathogens (**Fig. 1g**). While some pathogens exhibited maltose-dependent reduction in cocultures, none of the enterobacteria were suppressed entirely, unlike *A. baumannii* (**Fig. 1h**).

Together, these experiments demonstrate a significant reduction in *Acinetobacter* spp.-specific survival when both *K. oxytoca* and certain carbohydrates are present under anaerobic conditions.

### Suppression of *A. baumannii* by *K. oxytoca* occurs in hypoxic environments

Oxygen availability in the intestine varies significantly based on anatomical location, host health status, and microbial community composition, which are interconnected both directly and indirectly^58–62^. For example, broad-spectrum antibiotic treatment disrupts microbiota composition, resulting in higher oxygen concentrations in the intestine^62,63^. We next examined whether environmental oxygen conditions influence *A. baumannii* suppression. In coculture assays, complete suppression of *A. baumannii* was observed under anoxic and oxygen-limited conditions, such as incubation in an anaerobic chamber or using the AnaeroGen system, respectively (**Fig. 2a**). However, in cultures at atmospheric oxygen levels (∼21%), no suppression of *A. baumannii* was noted, despite comparable numbers of *K. oxytoca* recovered after 24 hours of incubation under all these conditions (**Supplementary Fig. 2a**). To identify oxygen concentrations that would enable suppression, coculture assays were performed under defined conditions in a hypoxic chamber (O_2_ conc.: 0%, 0.5%, 1%, 2%) (**Fig. 2b**). Suppression of *A. baumannii* was observed when oxygen levels were up to 1%. At 2% oxygen concentration, *A. baumannii* persisted, while *K. oxytoca* remained unaffected, maintaining consistent populations of approximately 10⁸ colony-forming units per milliliter (CFU/mL) (**Supplementary Fig. 2b**), underscoring its oxygen-independent growth potential.

**Fig. 2:**
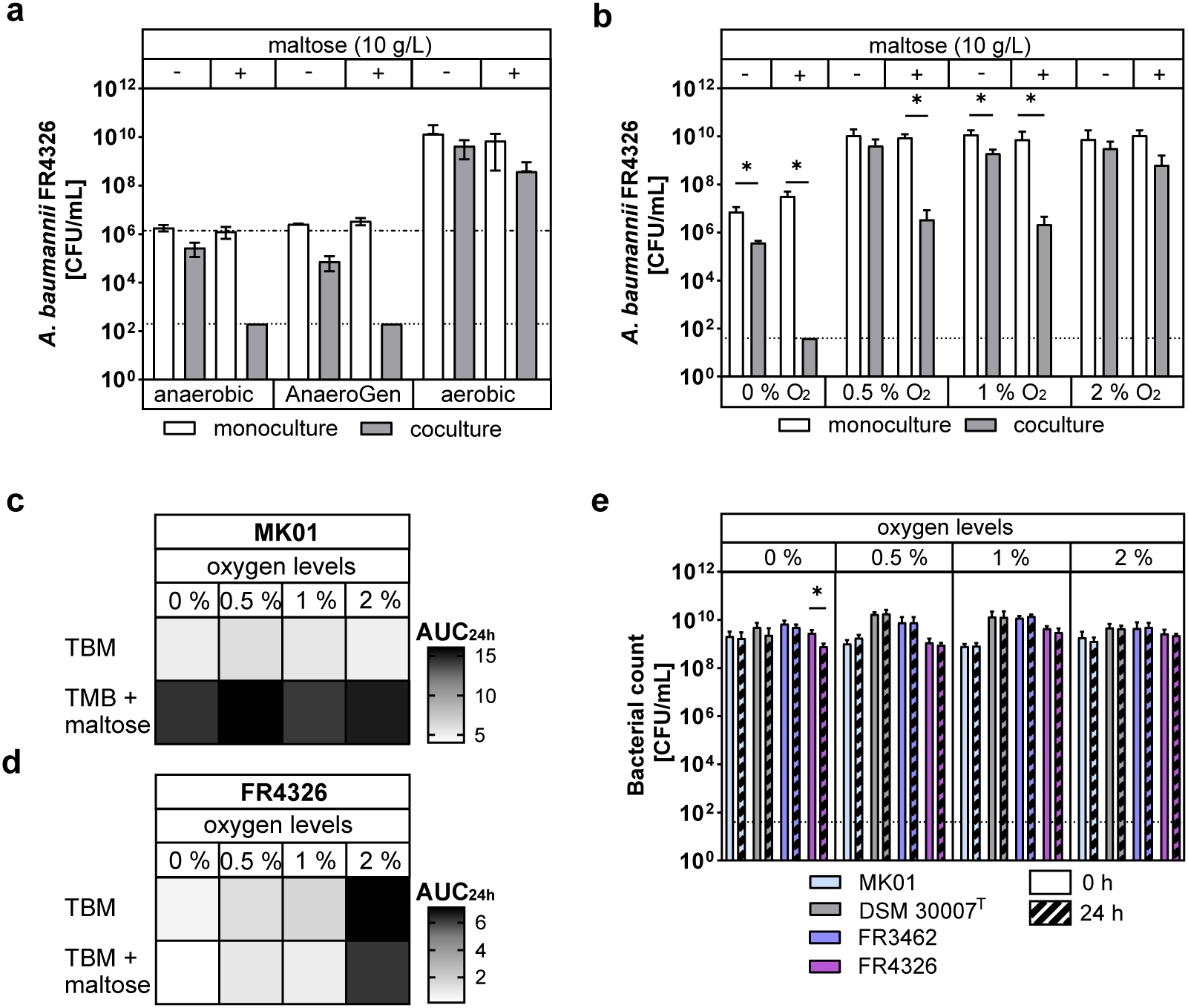
Effect of varying oxygen conditions and maltose on the growth of *A. baumannii* and *K. oxytoca* in monoculture and coculture. **a** *A. baumannii* FR4326 was cultured alone or in coculture with *K. oxytoca* MK01 in TBM ± 10 g/L maltose under different oxygen conditions: anaerobic chamber (0 % oxygen), AnaeroGen system, or aerobic conditions. The growth of *A. baumannii* was assessed through selective plating. Data are shown as mean ± SD for n= 3 biological replicates. **b** *A. baumannii* FR4326 was cultured alone or in coculture with *K. oxytoca* MK01 in TBM ± 10 g/L maltose under conditions with defined oxygen concentrations (0 % to 2 % oxygen). The growth of *A. baumannii* was assessed through selective plating. Data are shown as mean ± SD for n= 4 biological replicates. **c, d** *K. oxytoca* MK01 (**c**) and *A. baumannii* FR4326 (**d**) were grown in TBM ± 10 g/L maltose under conditions with defined oxygen concentrations (0 % to 2 % oxygen). Heatmaps of the Area Under the Curve (AUC) based on optical density OD_600_ measurements over 24 hours for n= 4 biological replicates. **e** Defined numbers of aerobically grown *K. oxytoca* MK01, *A. baumannii* DSM 30007^T^, FR3462 and FR4326 were plated on LB agar and incubated for 0 or 24 hours under hypoxic conditions (0 % to 2 % oxygen). Bacterial recovery was determined by CFU/mL with additional incubation information found in the methods. Data represent n= 4 biological replicates. The detection limit (dotted line) and the inoculum (mixed dashed line) are indicated. Statistical significance was determined using non-parametric Kruskal-Wallis test (**a**) and Mann-Whitney U test (**b**, **e**): ***p= 0.0006 (**a**) (*p < 0.05, **p<0.01, ***p<0.001).

Little is known about how oxygen availability itself influences *A. baumannii* growth and survival. Therefore, we first characterized its growth under defined oxygen concentrations and compared it to *K. oxytoca*. Under hypoxic conditions, *A. baumannii* exhibited distinct growth behavior, unlike *K. oxytoca*, which displayed robust growth. While *A. baumannii* formed colonies on solid media (**Fig. 2e**), its planktonic growth was severely restricted in liquid media when oxygen was limited (≤1 %) (**Fig. 2c, 2d; Supplementary Fig. 2c-j**). This suggests a limitation due to cell respiration and limited oxygen diffusion, which could affect metabolic processes, necessary for growth. This pattern was consistent across various *A. baumannii* strains (DSM 30007^T^, FR4326, and FR3462). Next, we assessed *A. baumannii* viability when exposed to varying oxygen concentrations but attached to a nutrient surface for up to 120 hours. Viability, based on recovered CFU, revealed no effect on *A. baumannii* survival under hypoxia after 24 hours at all tested oxygen concentrations (**Fig. 2e**). Notably, starting after 48 hours of oxygen exclusion, viable *Acinetobacter* recovery diminished in a strain-dependent manner (**Supplementary Fig. 2k**). Nevertheless, at oxygen levels above 0.5 %, *Acinetobacter* recovery remained stable at high numbers on agar plates (**Supplementary Fig 2l–n**). Together, these results reveal that active suppression of *A. baumannii* by *K. oxytoca* occurs at oxygen concentrations found in the intestinal environment.

### Metabolic by-/products secreted by *K. oxytoca*, rather than toxin production, lead to *A. baumannii* suppression

Both cell-contact-independent and -dependent mechanisms influence the outcomes of microbial interactions^64–66^. Consequently, coculture assays were conducted using a transwell setup to evaluate the role of cell-cell contact in *K. oxytoca*-mediated suppression of *A. baumannii*. Initially, monocultures of *K. oxytoca* and *A. baumannii* were separated by a 0.2 µm-membrane permeable only to metabolites and incubated anaerobically (**Fig. 3a**). Maltose-dependent inhibition of *A. baumannii* (approximately 10^4^-fold reduction) was observed, though full suppression was not achieved, even in the absence of cell contact (**Fig. 3c**). To determine whether slight differences in the experimental conditions between direct and transwell cocultures led to variations in the magnitude of suppression, we tested a second experimental setup. In this case, the two bacterial species were cocultured in one well, with the membrane separating the mixed cultures from an *A. baumannii* monoculture (**Fig. 3b**). When both bacterial species established cell-cell contact, *A. baumannii* experienced greater suppression in this setup (**Fig. 3d**). However, the suppression seemed to be contact-independent and did not require direct cell-contact for induction (**Fig. 3c and d**). These findings suggest that suppression is mainly mediated by a secreted factor produced by *K. oxytoca* under maltose-supplemented anaerobic conditions.

**Fig. 3:**
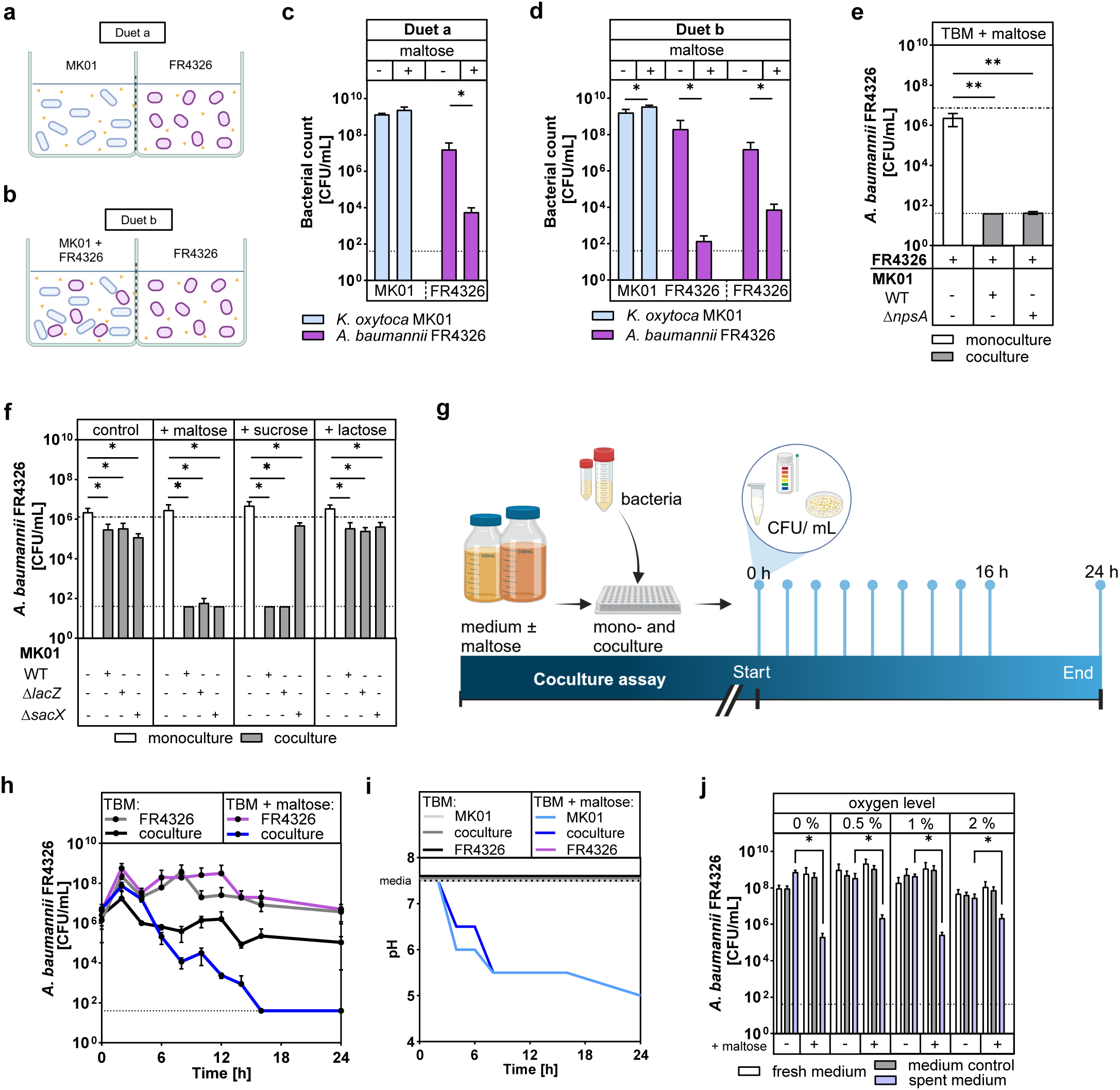
Suppression of *A. baumannii* is independent of cell-cell contact and *K. oxytoca* toxin production, but is driven by the metabolic activity of *K. oxytoca*. **a, b** Schematic display of configurations using the Cerillo Coculture Duet System with *K. oxytoca* MK01 (blue) and *A. baumannii* FR4326 (purple) cultured in mono- and cocultures separated by transwell membranes. **c, d** *K. oxytoca* MK01 (blue) and *A. baumannii* FR4326 (purple) were grown in monoculture and coculture in TBM ± 10 g/L maltose in membrane-separated wells in an anaerobic chamber. The growth of *A. baumannii* and *K. oxytoca* was assessed by selective plating. Data are shown as mean ± SD for n= 4 biological replicates. **e** *A. baumannii* FR4326 was grown alone or in coculture with *K. oxytoca* MK01 wt and *ΔnpsA* in TBM + 10 g/L maltose. The growth of *A. baumannii* was assessed by selective plating. Data are shown as mean ± SD for n= 3 biological replicates. **f** *A. baumannii* FR4326 was grown alone and in coculture with *K. oxytoca* MK01 wt, *ΔlacZ,* and *ΔsacX* in TBM ± 10 g/L maltose, sucrose or lactose. The growth of *A. baumannii* was assessed by selective plating. Data are shown as mean ± SD for n= 4 biological replicates. **g - i** Schematic workflow of longitudinal analysis of growth of *K. oxytoca* MK01 and *A. baumannii* FR4326 grown alone or in coculture in TBM + 10 g/L maltose. Parallel assays were stopped every 2 hours between 0 and 16 hours of incubation to collect supernatant for lactate quantification, growth enumeration (**h**), and pH measurement (**i**). The growth of *A. baumannii* (**h**) was assessed by selective plating. Data are presented as mean ± SD for n= 3 biological replicates. pH measurements (**i**) of the culture supernatants. j *K. oxytoca* MK01 was grown in TBM with and without maltose for 24 hours under hypoxic conditions (0 % to 2 % oxygen) in n= 4 biological replicates with incubation media controls. Steril-filtered spent media and controls, as well as fresh media were then used to culture initial OD_600_ 1 *A. baumannii* FR4326 under AnaeroGen conditions and plate on LB agar plates to display sole influence of supernatant on standardized cultures. Data are shown as mean ± SD for n= 4 biological replicates. All coculture assays were incubated under oxygen-limited conditions using the AnaeroGen system if not mentioned otherwise. The detection limit (dotted line) and the inoculum (mixed dashed line) are indicated. Statistical significance was determined using Mann-Whitney U test (**c**, **d**, **e**, **j**) and Wilcoxon matched-pairs signed rank test (**h,** **p = 0.0039) (*p < 0.05, **p<0.01).

*K. oxytoca* produces a genotoxin called tilimycin, whose procution is induced by various carbohydrates and exhibits antimicrobial activity against multiple bacterial species ^42,67^. To investigate whether suppression involves this toxin, we conducted coculture assays in tryptone maltose medium using wild-type *K. oxytoca* MK01 and a mutant deficient in toxin production, MK01 Δ*npsA* ^42^ (**Fig. 3e**). Coculture assays of *A. baumannii* with the wild-type and Δ*npsA* strains yielded comparable suppression, indicating that the suppression of *A. baumannii* is toxin-independent. To explore the relationship between suppression and carbohydrate utilization, we used *K. oxytoca* MK01 mutants, deficient in genes necessary for lactose (*lacZ*) and sucrose (*sacX*) utilization (**Fig. 3f; Supplementary Fig. 3a**). Suppression of *A. baumannii* was maintained by all strains in maltose-containing medium, while lactose supplementation had no significant impact as previously observed (**Fig. 1d**). In a sucrose-supplemented medium, *A. baumannii* was suppressed by both wild-type and *lacZ* mutant strains but not by the *sacX* mutant, highlighting that the ability to utilize a specific carbohydrate is essential for suppression.

To identify when *A. baumannii* suppression occurs during coculture, we longitudinally quantified *A. baumannii* survival, accompanied by pH measurements as an indirect readout of *K. oxytoca*’s anaerobic metabolism (**Fig. 3g**). A rapid decline in *A. baumannii* survival was detectable within the first eight hours, and no colonies could be recovered after 16 and 24 hours (**Fig. 3h**). The decline in *A. baumannii* survival correlated with a significant drop in pH from 7.5 to 5.5, indicating active *K. oxytoca* metabolism (**Fig. 3i; Supplementary Fig. 3b**). Finally, to corroborate the involvement of secreted metabolites or metabolic byproducts, supernatants from 24-hour *K. oxytoca* monocultures were generated under varying oxygen concentrations and in the presence or absence of maltose, sterile filtered, and then used to incubate defined amounts of *A. baumannii* for another 24 hours (**Fig. 3j; Supplementary Fig. 3c**). Only the supernatant from maltose-containing *K. oxytoca* cultures grown under hypoxic conditions led to a significant (>10^3^-fold) reduction of *A. baumannii* survival. This further supports the hypothesis that inhibitory factors are secreted under oxygen-limiting, maltose-metabolizing conditions.

### Selective metabolites impact the competitive dynamics of *K. oxytoca* and *A. baumannii*

In contrast to aerobic growth, the anaerobic growth of *K. oxytoca* leads to the acidification of the culture media due to carbohydrate fermentation. To evaluate whether acidification itself contributed to the suppression of *A. baumannii*, monocultures of *A. baumannii* and *K. oxytoca,* as a control, were incubated in TMB that was adjusted to initial pH values ranging from 3.5 to 9.5. After 24 hours, bacterial survival and growth, respectively, were quantified (**Fig. 4a**). *A. baumannii* persisted at neutral to alkaline pH, showing a tenfold reduction in recovery at acidic pH (5.0–5.5). These findings suggest that acidification may contribute to the suppression of *A. baumannii*. Next, coculture assays were performed in media buffered to acidic (pH 5), neutral (pH 7.5), and alkaline (pH 9) conditions using MES, MOPS, or TAPS buffers (**Supplementary Fig. 4a -4d**). In cocultures with *K. oxytoca*, suppression occurred across all buffered conditions, with *A. baumannii* CFU counts reduced to the detection limit at lower buffer concentrations (50 mM). At higher buffer concentrations (200 mM), *A. baumannii* survival improved modestly in neutral and alkaline conditions (10³–10⁴ CFU/mL). Control experiments demonstrated that survival and growth in buffered media were not significantly impaired for monocultures and cocultures without maltose (**Supplementary Fig. 4a, 4b**). These results indicate that while acidification contributes to the observed suppression, it acts synergistically with other mechanisms mediated by *K. oxytoca*.

**Fig. 4:**
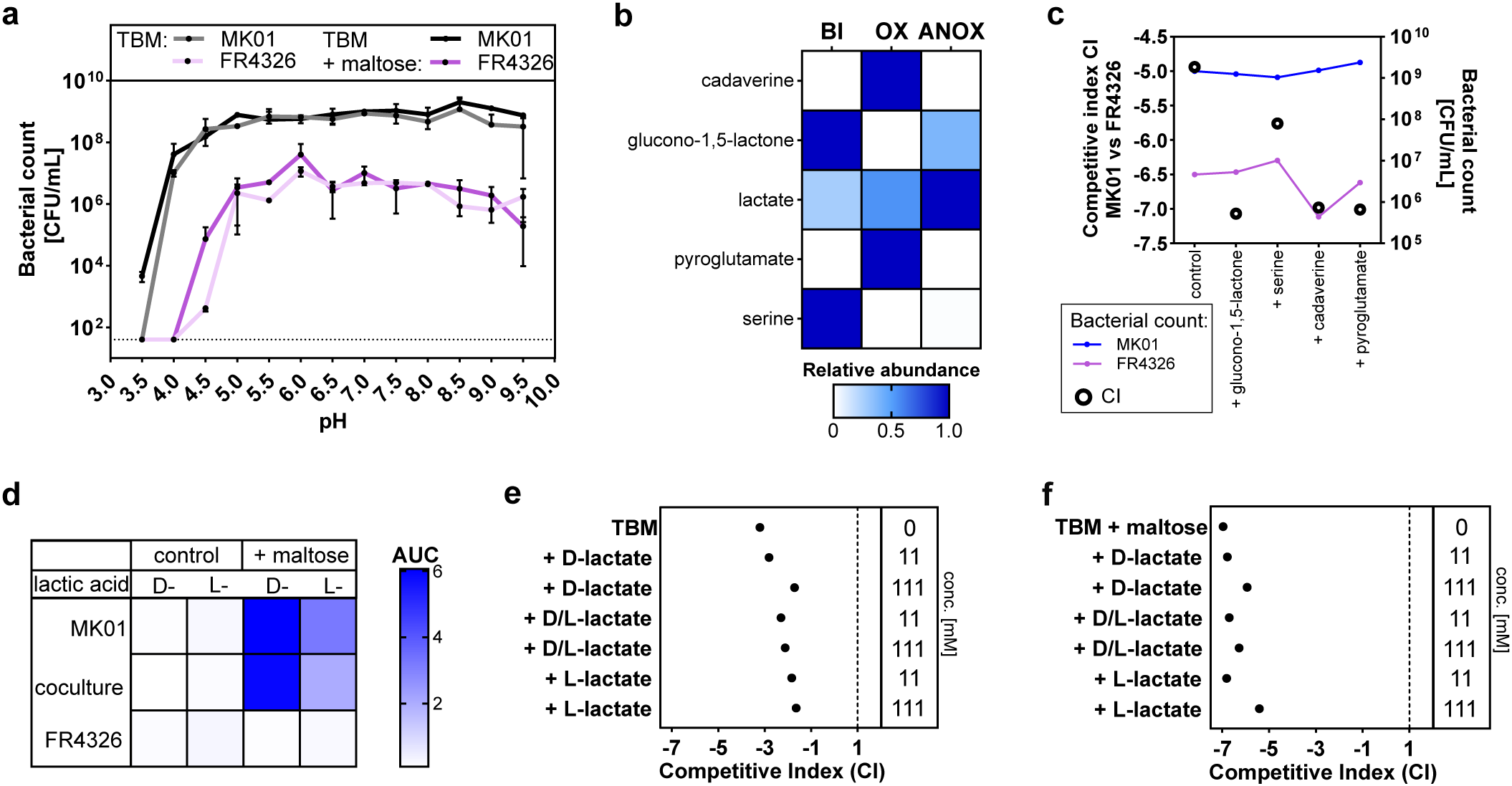
Environmental and metabolic influences on *K. oxytoca* and *A. baumannii* growth and competition. **a** *K. oxytoca* MK01 and *A. baumannii* FR4326 were cultured in TBM ±10 g/L maltose in initial media pH from 3.5 to 9.5. The growth of standardized bacterial inocula for both bacteria was assessed by plating after 24 hour incubation on LB agar plates and the detection limit (dotted line) is indicated. Data are shown as mean ± SD for n= 2 biological replicates. **b** GC-MS metabolomic analysis of *K. oxytoca* MK01 culture supernatant in TBM + 10 g/L maltose. Samples were collected before incubation (BI), and after 24 hours of incubation under aerobic (OX) or anaerobic (ANOX) conditions in an anaerobic chamber. Relative abundances of selected detected compounds are shown, normalized to each compound itself. **c** Compositional figure to display competitive indices (CI) and bacterial growth of *K. oxytoca* MK01 and *A. baumannii* FR4326 after mono- and coculture in TBM with 10 g/L maltose and the addition of either 9.8 mM cadaverine, 5.6 mM glucono-1,5-lactone, 7.7 mM pyroglutamate or 38 mM serine. Data are shown as mean ± SD for n= 3 biological replicates. **d** Heatmap displaying the Area under the curve (AUC) of lactic acid accumulation in *K. oxytoca* MK01-culture supernatant over time, as measured by enzymatic detection. Supernatant was collected during longitudinal growth analysis of *K. oxytoca* MK01 and *A. baumannii* FR4326 mono- and cocultures in n = 3 biological replicates (Fig. 3g). **e, f** Competitive indices (CI) of *K. oxytoca* MK01 and *A. baumannii* FR4326 mono- and cocultures in TBM (**e**) or TBM + 10 g/L maltose (**f**) with the addition of lactate. Sodium -D-, - D/L or -L-lactate was added to lactate concentrations of 11 mM or 111 mM. Data are shown as mean ± SD for n= 3 biological replicates. All mono- and coculture assays were incubated under oxygen-limited conditions using the AnaeroGen system if not mentioned otherwise.

To identify metabolites associated with *A. baumannii* suppression, we hypothesized that comparing the growth media from aerobic and anaerobic *K. oxytoca* cultivation-representing non-suppressing and suppressing potential-could serve as a starting point for further analyses. Notably, metabolomic analysis of *K. oxytoca* supernatants using gas chromatography-mass spectrometry revealed distinct differences between oxygen-rich and anoxic conditions (**Fig. 4b, Supplementary Fig. 4e**). Specifically, the analysis identified various compounds produced exclusively in aerobically incubated cultures, in anaerobically incubated cultures, or remaining present in the environment after exposure to either oxygen condition. For instance, high concentrations of lactate were detected in anaerobic cultures. Additionally, cadaverine and pyroglutamate were found in high concentrations in supernatants from aerobically grown cultures, alongside significant depletion of glucono-1,5-lactone and serine, which were initially present in the medium but largely disappeared after anaerobic growth. These findings suggest that nutrient competition or selective metabolite production could influence *A. baumannii* survival in specific environments. However, since *A. baumannii* suppression occurs only under anoxic conditions, the mechanism likely involves metabolites produced or retained specifically in oxygen-limited environments.

To directly investigate the role of metabolite candidates in *K. oxytoca-A. baumannii* interactions (**Fig. 4b**), we performed coculture assays in TBM with or without maltose supplementation, adding the candidates at varying concentrations. The addition of cadaverine (0 - 98 mM) (**Supplementary Fig. 4f**) or pyroglutamate (0 – 77 mM) (**Supplementary Fig. 4h**), two compounds negatively correlated to *A. baumannii* suppression, did not prevent *A. baumannii* suppression by *K. oxytoca* or growth in monocultures or cocultures at lower concentrations. However, at the highest concentrations, both compounds exhibited lethal effects on *A. baumannii* in both monoculture and coculture. Supplementation of glucono-1,5- lactone (0 – 56 mM) (**Supplementary Fig. 4g**), which is enriched under anaerobic conditions, to TBM had no significant impact on *A. baumannii* suppression or growth at lower concentrations. Notably, the compound was lethal to *A. baumannii* at the highest tested concentration. In coculture assays with maltose supplementation, the addition of cadaverine, pyroglutamate, or glucono-1,5-lactone at sublethal concentrations failed to reverse *A. baumannii* suppression (**Fig. 4c; Supplementary Fig. 4k**). Serine was depleted by *K. oxytoca* under aerobic conditions and, to a lesser extent, under anaerobic incubation in maltose-containing medium (**Fig. 4b**). Reportedly, serine impacts bacterial virulence and stress response^68–72^, and specifically *A. baumannii* survival in complex environments^73^, potentially mitigating adverse environmental conditions. To test its effect on *A. baumannii*, serine (0, 19 mM, 38 mM) was added to TBM at pH values of 5, 6, and 7.5, followed by *A. baumannii* monoculture incubation in these media (**Supplementary Fig. 4i**). Moderate increases in *A. baumannii* CFU/mL were observed at pH 6 and 7.5 with lower serine concentrations, but no significant impact was evident at acidic pH or higher concentrations. In coculture assays with maltose-supplemented medium and 38 mM serine, *A. baumannii* suppression was unaffected (**Supplementary Fig. 4j**). Under acidic starting conditions (pH 5), *A. baumannii* monoculture CFU/mL decreased further, highlighting limited support from serine in competitive dynamics. The final tested metabolite, lactate, was detected in the supernatants of *K. oxytoca* monocultures and cocultures with *A. baumannii*, but not in *A. baumannii* monoculture supernatants (**Fig. 4d; Supplementary Fig. 4l-n**). Differentiating lactate enantiomers, predominantly D-lactate, was identified as a product of *K. oxytoca* metabolism. To evaluate its impact on the bacterial interaction, we added lactate (in L-, D-, and racemic forms) at increasing concentrations to TBM without maltose (**Fig. 4e; Supplementary Fig. 4o, 4p**). In this setup, higher lactate concentrations reduced *K. oxytoca* dominance in cocultures to a noticeable degree, with *A. baumannii* coculture CFU/mL comparable to those observed in respective *A. baumannii* monocultures. When lactate was added to tryptone medium with maltose (**Fig. 4f; Supplementary Fig. 4q, 4r**), the competitive index revealed a stronger *K. oxytoca* dominance in coculture interactions. *A. baumannii* monoculture CFU/mL were noticeably lower at higher lactate concentrations, and *A. baumannii* reduction in coculture to the detection limit of the assay was seen across all tested lactate enantiomers and the racemic mixture (**Supplementary Fig. 4q**). Interestingly, at higher lactate concentrations, small populations of *A. baumannii* (∼ 10^2^ CFU/mL) were recoverable from cocultures, while *K. oxytoca* consistently maintained recoverable high cell densities (∼ 10^8^ CFU/mL).

These results highlight that neither the presence of lactate nor any other of the tested metabolites interfered with the outcome of *K. oxytoca*-mediated *A. baumannii* suppression or were able to induce the same suppression themselves. Future studies could explore synergistic effects with additional metabolites and/or an acidic environment.

### Carbohydrate utilization driven suppression of *A. baumannii* by *K. oxytoca* and commensal microbes in ex vivo and *in vivo* gut models

Since we could not identify a specific factor responsible for the maltose-dependent suppression of *A. baumannii* by *K. oxytoca*, we aimed to characterize whether this effect is a common trait among members of the KoSC. Therefore, we conducted coculture assays using strains of *K. oxytoca*, *K. grimontii*, and *K. michiganensis* at a 10:1 ratio of *Klebsiella* spp. to *A. baumannii* in TBM, with and without maltose supplementation (**Fig. 5a**). Across all tested strains, *A. baumannii* was consistently reduced in a maltose-dependent manner to or near the detection limit, suggesting that suppression is a shared feature of the KoSC. To assess whether this effect extends to other commensal intestinal bacteria, we tested a panel of *Citrobacter braakii, Enterobacter ludwigii, Enterococcus faecium, Lactobacillus paracasei, Lactococcus lactis,* and *Streptococcus oralis* in modified Gifu Anaerobic Medium (mGAM), with and without maltose addition (**Fig. 5b, Supplementary Fig. 5a**). In maltose-supplemented mGAM, *K. oxytoca* reduced *A. baumannii* to 10³ CFU/mL with *E. faecium* and *S. oralis* demonstrating similar suppression of the pathogen, while most other commensal bacteria were unable to achieve the same effect.

**Fig. 5:**
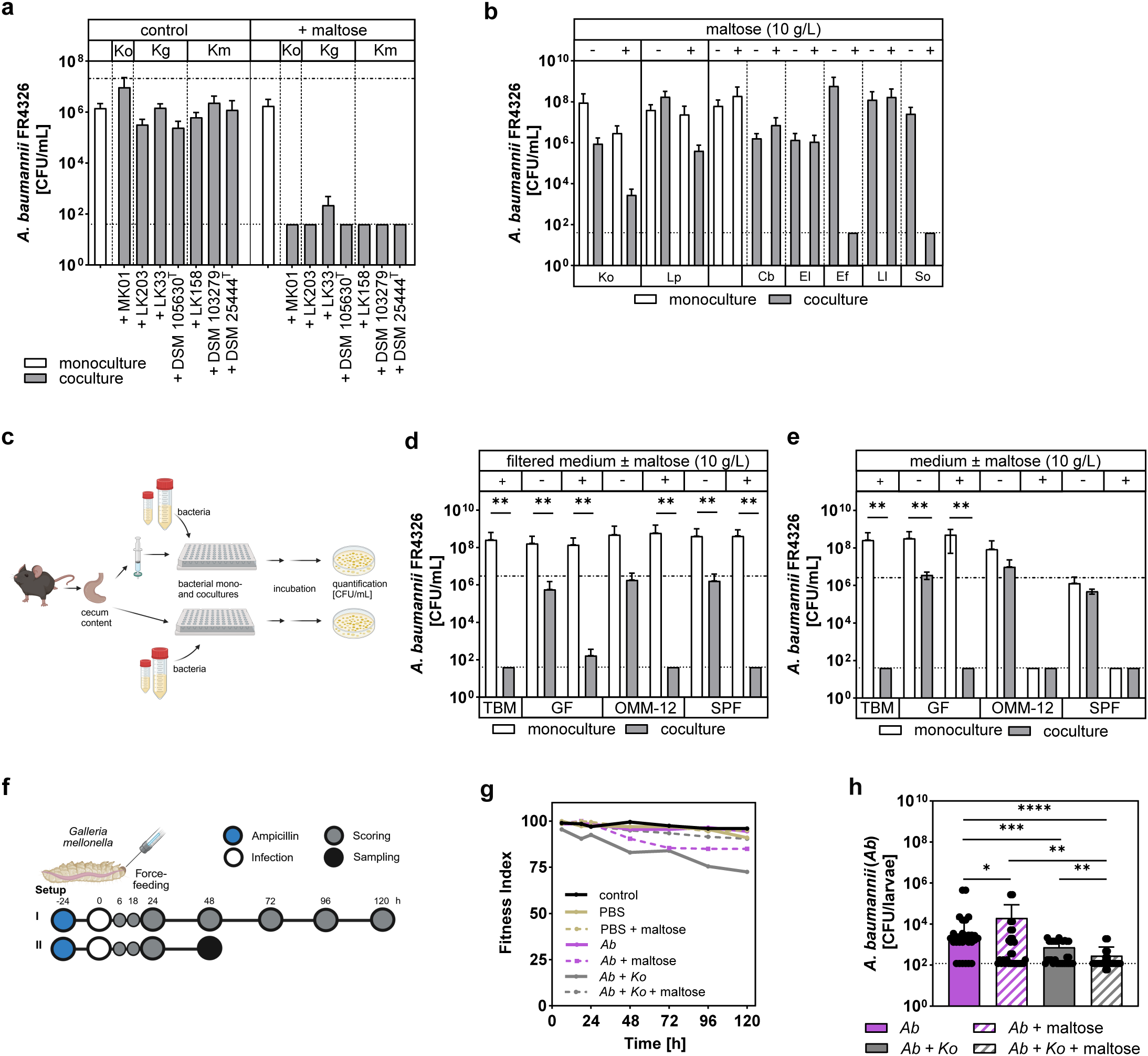
Carbohydrate utilization driven suppression of *A. baumannii* in various intestinal models. **a** *A. baumannii* FR4326 was grown alone and in coculture with *Klebsiella oxytoca* species complex (KoSC) strains in TBM ± 10 g/L maltose. The growth of *A. baumannii* was assessed through selective plating. The mean ± SD for n= 2 biological replicates is displayed. Included strains are *K. oxytoca* (Ko) MK01*, K. grimontii* (Kg) LK203, LK233 and DSM 105630^T^, and *K. michiganensis* (Km) LK158, DSM 103279 and DSM 25444^T^. **b** *A. baumannii* FR4326 was grown alone and in coculture with various intestinal commensal species in mGAM ± 10 g/L maltose. The growth of *A. baumannii* was assessed through selective plating. The mean ± SD for n= 3 biological replicates is displayed. Commensal strains are isolates from healthy human donors: *K. oxytoca* MK01 (Ko), *Lactobacillus paracasei* LK 355 Iso 2 (Lp), *Citrobacter braakii* LK 280 Iso 12 (Cb), *Enterobacter ludwigii* LK 290 Iso 14 (El), *Enterococcus faecium* LK 290 Iso 7 (Ef), *Lactococcus lactis* LK 370 Iso 1 (Ll), *Streptococcus oralis* LK 323 Iso 5 (So). **c - e** Schematic workflow of a coculture assay using murine cecum content as the incubation medium to grow *A. baumannii* FR4326 alone and in coculture with *K. oxytoca* MK01 with and without 10 g/L maltose. Germ-free (GF), specific-pathogen-free (SPF) or OMM-12 – carrying cecum content was used either filtered (**d**) or unfiltered (**e**) in a 1:20 dilution in phosphate-buffered saline (PBS). The growth of *A. baumannii* was assessed through selective plating. Data show the means ± SD for n= 6 biological replicates of bacteria, cultured in n= 2 biological replicates of cecum content. **f – h** Schematic workflow of a *Galleria mellonella* model, where larvae were force-fed with either *A. baumannii* FR4326 (*Ab*) alone or in combination with *K. oxytoca* (*Ab + Ko*), with or without maltose supplementation. (**g**) Larval fitness of control groups and after bacterial administration was assessed according to an adapted Soring model for n= 2 x 10 larvae (Setup I). (**h**) Bacterial recovery from larval homogenates was determined 48 h post-infection by selective plating (Setup II). Data represent mean ± SD from at least 27 larvae per condition (two independent bacterial culture replicates). Pairwise statistical differences were analyzed using the Mann–Whitney U test (****p < 0.0001). Overall statistical significance was determined by Kruskal–Wallis test (****p < 0.0001). All coculture assays were incubated under oxygen-limited conditions using the AnaeroGen system if not mentioned otherwise. The detection limit (dotted line) and the inoculum (mixed dashed line) are indicated. Statistical significance was determined using Mann-Whitney U test (*p < 0.05, **p<0.01).

To investigate whether the suppressive effect of *K. oxytoca* would be observable under the nutrient conditions of the gut, coculture assays were conducted using cecal content from mice with distinct gut microbiota profiles: specifically, germ-free, gnotobiotic OMM-12 ^74^, and specific pathogen-free (SPF) mice under oxygen-limited conditions (**Fig. 5c**). We utilized sterile-filtered cecal content diluted 1:20 in phosphate-buffered saline (PBS) as a bacterial culture medium that represents a complex nutrient environment with the dilution allowing for efficient use of cecum content (**Fig. 5d; Supplementary Fig. 5b**). Under these conditions, *A. baumannii* grew efficiently, and maltose supplementation alone did not affect *A. baumannii* monoculture growth. In coculture scenarios, *A. baumannii* was significantly suppressed by *K. oxytoca* in a maltose-dependent manner. Subsequently, we performed coculture assays using unfiltered cecal content (1:20 dilution in PBS) to account for the inherent microbial activity (**Fig. 5e; Supplementary Fig. 5c**). In the absence of maltose, *A. baumannii* could colonize all microbial environments, but growth varied by 10^2^-fold, being lowest in the presence of the complex SPF microbiota. Maltose supplementation completely suppressed *A. baumannii* in both the OMM-12 and SPF microbiota communities, which contain endogenous *Enterococcus* strains, even without *K. oxytoca*. These findings suggest that besides *K. oxytoca*, other inherent microbiota members may be stimulated by maltose to inhibit *A. baumannii* in complex microbial settings.

To further assess the in vivo relevance of these observations, we used Galleria mellonella as a model of intestinal infection and colonisation ^75–77^. Although *G. mellonella* is frequently used in *A. baumannii* research, oral administration via force-feeding is rarely employed, particularly in studies focusing on gut colonisation and carbohydrate-mediated microbial interactions ^53,78^. Nevertheless, force-feeding ensures a defined bacterial input, prevents environmental contamination and enables control at the level of the individual larva. Of note, force-feeding of neither *A. baumannii* and *K. oxytoca* caused no significant changes in fitness or survival across groups (**Fig. 5g**, **Supplementary Fig. 5d, 5e, 5g**), as assessed by an adapted survival scoring considering survival, activity, melanization and cocoon/pupa formation ^76,83^.

First, the colonization potential of was evaluated (**Supplementary Fig. 5f**). The larvae were either left untreated or pretreated with ampicillin 24 hours before infection, in order to reduce the native microbiota and increase the size of the intestinal niche.. Each larva received 10^7^ CFU of *A. baumannii* or *K. oxytoca* per feeding. At 48 h post-infection, recoverable *A. baumannii* was markedly higher in ampicillin-pretreated larvae compared to untreated controls, consistent with increased niche availability. Snap-freezing and resection of larvae allowed exclusion of non-gut tissue colonization, confirming that recovery reflected true gastrointestinal colonization. Next, we examined the effect of carbohydrate supplementation (**Fig. 5f**). When maltose was added, a slight increase in recoverable separately force-fed *A. baumannii* and *K. oxytoca* was observed, suggesting improved colonization under carbohydrate-enriched conditions (**Fig. 5h, Supplementary Fig. 5h**). However, the most pronounced suppression of *A. baumannii* occurred when it was co-administered with *K. oxytoca* in the presence of maltose. Under these conditions, *A. baumannii* recovery dropped significantly, while *K. oxytoca* proliferated and exceeded *A. baumannii* CFU per larva (**Supplementary Fig. 5h**).

These results demonstrate that the *Galleria* model enables controlled intestinal colonization experiments, precise carbohydrate supplementation, and stepwise translation along the *in vitro*–*ex vivo*–*in vivo* axis. They also confirm that the combination of *K. oxytoca* and maltose exerts the strongest suppressive effect on *A. baumannii in vivo*.

## Discussion

The findings of this study provide insights into the competitive interactions between *K. oxytoca* and *A. baumannii*, whose outcome is regulated by the absence of oxygen and carbohydrate availability. Our results suggest that commensal gut bacteria, such as *K. oxytoca,* can suppress *A. baumannii* in environments supplemented with specific carbohydrates, like maltose. This suppression is mediated by secreted metabolites and the acidification of the surrounding environment without relying solely on direct cell-cell contact among the bacterial members. Importantly, this effect was not limited to *ex vivo* culture systems: *in vivo* validation in ampicillin-pretreated *Galleria mellonella* larvae demonstrated that recoverable *A. baumannii* was significantly reduced when larvae were co-force fed with *K. oxytoca* and maltose, confirming the robustness of this suppressive interaction under host-associated conditions.

*A. baumannii* is a clinically relevant opportunistic pathogen, with systemic infections often originating from intestinal colonization. The intestinal tract serves as a critical reservoir for *A. baumannii*, particularly in neonates, immunocompromised individuals, or patients undergoing prolonged antibiotic therapy ^48,84^. Strategies to reduce intestinal reservoirs of *A. baumannii* could, therefore, play a crucial role in preventing systemic infections and, especially, the even more severe *A. baumannii* coinfections with other potent human pathogens, as well as in controlling nosocomial outbreaks ^85,86^. *A. baumannii* is reported as an aerobic bacterium; however, it can be found in both oxic and typically anoxc environments ^87–90^, but also hypoxic environments such as the intestinal tract of animals and humans ^61,91–94^. Our findings suggest it can persist under anoxic conditions and even grow under hypoxic conditions. Under strictly anoxic conditions, *A. baumannii* can survive for extended periods on surfaces, though its growth is limited. At oxygen levels as low as 0.5 %, the decline of *A. baumannii* on surfaces diminishes, and at 2 % oxygen, bacterial growth is restored in liquid media, demonstrating its ability to adapt and thrive in various environments. This adaptive capacity underscores the importance of targeting *A. baumannii* even within intestinal niches with low oxygen availability.

Within this context, the gut microbiome can serve as a natural defense against *A. baumannii*. Our *Galleria mellonella* experiments demonstrate that deliberate supplementation with specific carbohydrates, such as maltose, can enhance the fitness of probiotic strains like *K. oxytoca* and shape the intestinal microbiome in a favorable manner. In this model, maltose supplementation and *K. oxytoca* treatment synergistically suppressed *A. baumannii* colonization in vivo, highlighting the importance of targeted metabolic interactions. Force-feeding in larvae enabled precise control of bacterial and carbohydrate input, minimized confounding effects from dietary intake, and maintained larval fitness, illustrating the utility of Galleria as a tractable and ethically advantageous system for bridging *in vitro*, *ex vivo*, and *in vivo* studies.

These findings suggest that combining probiotic administration with selective carbohydrate supplementation could complement current antimicrobial strategies by transiently fortifying the gut microbiome and reducing intestinal *A. baumannii* reservoirs without reliance on traditional antibiotics. Probiotic formulations containing *K. oxytoca* together with defined carbohydrates may therefore represent a novel strategy to control pathogen colonization and limit the risk of systemic dissemination.

Future studies should extend these observations to mammalian models to evaluate the efficacy of carbohydrate-probiotic interventions in more complex gut environments, where host immunity, microbial diversity, and dietary intake may modulate colonization dynamics. Controlled experiments in mammals would allow systematic assessment of carbohydrate availability, probiotic dosing and safety ^95,96^, and timing to optimize suppression of *A. baumannii*. Furthermore, mechanistic investigations in these systems could elucidate how *K. oxytoca* competes with pathogens and how specific metabolites mediate colonization resistance. Collectively, such studies would provide a strong preclinical foundation for designing translational interventions aimed at preventing gut colonization and systemic spread of multidrug-resistant *A. baumannii*.

In conclusion, our study underscores the central role of carbohydrate metabolism in shaping competitive interactions between gut-associated pathogens and commensals. Temporarily fortifying the microbiome through selective carbohydrate supplementation and probiotic administration emerges as a promising, safe, and mechanistically informed strategy to reduce intestinal *A. baumannii* reservoirs and mitigate the risk of systemic infections.

## Material and Methods

### Contact for reagent and resource sharing

Additional information, as well as requests for resources and reagents, should be directed to the lead Contact, Prof. Dr. Till Strowig.

### Ethics statement

All experiments working with substances of animal origin were performed in agreement with the guidelines of the Helmholtz Center for Infection Research, Braunschweig, Germany, the National Animal Protection Law (Tierschutzgesetz (TierSchG), the animal experiment regulations (Tierschutz-Versuchstierverordnung (TierSchVersV)), and the recommendations of the Federation of European Laboratory Animal Science Association (FELASA).

### Mice

Germ-free (GF) C57BL/6NTac mice and C57BL/6N mice with OMM-12 microbiome were bred in isolators in the Germ-free facility at the Helmholtz Center for Infection Research (HZI) animal facility on campus. C57BL/6N SPF mice were bred and maintained at the HZI animal facility on campus. All mice were killed by euthanization with CO_2_ and cervical dislocation. Subsequently, cecum content for experiments was collected.

### Bacterial strains

The reference strain *K. oxytoca* MK01 originated from stool of a human donor. Further information can be found in previous studies^40,42^. Deletion mutants Δ*npsA* and Δ*sacX* were generated and used in previously published work ^41,42,97^. The deletion mutant Δ*lacZ* was generated according to a previously published protocol^97^. Strains are available upon request from the lead contact.

*A. baumannii* FR4326 and *A. baumannii* FR3462 were isolated from patients and provided by Volkhard A. J. Kempf from the University Hospital Frankfurt.

Additional bacterial strains for screening purposes were obtained from cooperation partners at: Hannover Medical School (MHH), the National Reference Laboratory for multidrug-resistant gram-negative bacteria (NRZ), the Leibniz Institute DSMZ-German Collection of Microorganisms and Cell Cultures GmbH, through its DZIF Pathogen Repository and from the University Hospital Magdeburg (OVGU), and Humboldt University Berlin.

Intestinal commensal bacterial strains were isolated from stool of healthy human donors, participating in the LöwenKIDS Cohort (Clinicaltrials.Gov Identifier: NCT02654210) of the Martin-Luther-University Halle-Wittenberg. Detailed information on the isolation procedure can be found in the method section under strain isolation. Further information on the human cohort can be found in the published cohort profile^98^.

Details on all used bacterial strains can be found in the bacterial strain list (**Supplementary Table 1**).

### Biolog Phenotype MicroArray Assay

Metabolic profiling of strains was performed using the Biolog Phenotype MicroArray system. Bacterial strains were cultivated on R2A agar plates (Difco) overnight. Bacterial material was collected and resuspended in phosphate-buffered saline (PBS) to an optical density at 600 nm (OD_600_) of 0.2. For each well of the PM1 and PM2A Biolog microplates, 99 µL of saline Minimal Medium (MM9) without carbon sources and 1 µL of bacterial suspension was added. Cultures were incubated for 24 hours aerobically at 37 °C in a microplate spectrophotometer (BioTek LogPhase 600 Microbiology Reader) with continuous shaking and bacterial growth was monitored by measuring OD_600_ every hour. Only substrates resulting in OD_600_ values above 0.2 were considered for analysis. Growth patterns and carbon source utilization were evaluated by calculating the area under the curve (AUC).

### *In vitro* coculture Assay

Coculture assays were conducted to evaluate bacterial competition in minimal isotonic tryptone medium with optional carbohydrate supplementation. Standardized coculture assays were performed in 96-well plates (TC-Plate Standard F with lid, Sarstedt) using a 10:1 initial inoculation ratio of *K. oxytoca* to *A. baumannii*, simulating a competitive environment with an established commensal community. The incubation tryptone base medium (TBM) consisted of 17 g/L tryptone/ peptone (Carl Roth) with the addition of 2.5 g/L K_2_HPO_4_ (Merck) and an optional addition of 10 g/L maltose (Sigma-Aldrich). Bacterial inocula were prepared from aerobic overnight cultures in LB medium (Carl Roth). Well plates were incubated at 37 °C with shaking under oxygen-limited conditions, installed with an AnaeroGen system in a sealed box. Oxygen-exclusion was monitored routinely with Anaerotest strips (Millipore). An input aliquot was diluted in PBS and plated for CFU/mL estimations and culture contaminations on respective agar plates. After 24 hours, cultures were serially diluted in PBS and 25 µL of each dilution was spread with a sterile inoculation loop on a quarter of a selective agar plate to differentiate and quantify bacterial species (*A. baumannii* on LB agar (Carl Roth) + antibiotic; *K. oxytoca* on CHROMagar orientation (CHROMagar). Plates were incubated at 37 °C for 24 hours with subsequent bacterial colonies counting and colony-forming units (CFU/mL) calculations.

Adaptations of this standardized coculture assay include:

#### a) Carbohydrate screening

To assess the utilization and impact of various carbohydrate, 10 g/L of selected saccharides were added to tryptone base medium. The supplementary compounds were chosen according to Biolog results. The following saccharides were used: Fructose (Carl Roth), D-(+)-galactose (Sigma-Aldrich), D-(+)-glucose (Sigma-Aldrich), D-(+)-mannose (Sigma-Aldrich), L-sorbose (Carl Roth), D-(+)-cellobiose (Sigma-Aldrich), isomaltose (Sigma-Aldrich), α-lactose (Sigma-Aldrich), lactulose (Carl Roth), D-(+)-maltose (Sigma-Aldrich), D-(+)-sucrose (Sigma-Aldrich), D-(+)-trehalose (Carl Roth), maltotriose (Sigma-Aldrich), D-(+)-raffinose (Carl Roth), (+)-arabinogalactan from larch wood (Sigma-Aldrich), starch (Sigma-Aldrich), xylan from corn (abcr). Supplemented media were sterile-filtered using syringes (Sarstedt) and filters (Filtropur S, Sarstedt). Coculture assays according to the standardized protocol were conducted in multiple sets and rounds of testing. Displayed is the composition of tested carbohydrates with assay controls, tryptone base medium without carbon sources and with maltose addition for every test set.

#### b) Maltose-supplementation parameters

Exemplarily for carbohydrate supplementation, different concentrations of maltose addition to tryptone base medium without a carbohydrate were tested in the standardized coculture assay. Maltose concentrations ranged from 0 g/L to 20 g/L. To eliminate tryptone base medium as most influential in the coculture results, maltose supplementation of other media was tested. Incubation media used in the standardized coculture assay in addition to tryptone base medium with maltose were saline Minimal Medium (MM9), LB medium (Carl Roth) and modified Gifu Anaerobic Medium (mGAM, himedia). Every additional medium was tested with and without 10 g/L maltose supplementation.

#### c) Buffer supplementation

To stabilize the pH environment during bacterial coculture without directly influencing the involved bacteria, Good’s buffer were added to tryptone base medium with or without 10 g/L maltose supplementation. To cover acidic, neutral and alkaline conditions, the following buffer were used: 2-morpholin-4-ylethanesulfonic acid (MES, Carl Roth) adjusted to pH 5, 3-(morpholin- 4-yl)propane-1-sulfonic acid (MOPS, Carl Roth) for pH 7 and 3-{[1,3-Dihydroxy- 2- (hydroxymethyl)propan-2-yl]amino}propane-1-sulfonic acid (TAPS, Sigma-Aldrich) for pH 9. Buffers were added to the medium with installed pH using a pH meter (Mettler Toledo) at pH 5, pH 7 and pH 9 at final concentrations of 50 mM or 200 mM prior to bacterial inoculation. Subsequently, the standardized coculture assay was performed. Instead of plating diluted cultures after incubation, 5 µL droplets of each serial dilution were placed on selective agar plates. Plates were incubated with agar side down at 37 °C for 24 hours. After incubation, bacterial colonies were counted in each droplet and CFU/mL calculated with a respectively higher detection limit.

#### d) Addition of selected compounds

From metabolomic analysis, five compounds were selected to investigate further hypotheses regarding *A. baumannii* suppression in *K. oxytoca* coculture using the standardized coculture assay: cadaverine (Sigma-Aldrich) and D/L-pyroglutamate (Sigma-Aldrich), D-(+)-glucono-1,5-lactone (Thermo Scientific) and L-serine (Carl Roth) and sodium L-lactate (Sigma-Aldrich), sodium D-lactate (Sigma-Aldrich) and sodium D/L-lactate (Sigma-Aldrich). Supplement concentrations were informed estimates based on literature research. Cadaverine was added to the tryptone base medium to final concentrations of 0.98, 9.8 or 98 mM^99,100^ as well as D/L-pyroglutamate at concentrations of 0.77, 7.7 and 77 mM. D-(+)-glucono-1,5-lactone was added at 0.56, 5.6 and 56 mM ^101,102^, while L-serine was tested at final concentrations of 19 or 38 mM ^68,73,103^. Following 24 hours of incubation, all cultures were processed according to the standard protocol under AnaeroGen conditions.

#### e) Ratio of *K. oxytoca* to *A. baumannii*

To assess the influence of competitor ratios, standardized coculture assays were conducted using initial inoculation ratios of *K. oxytoca* to *A. baumannii*, ranging from 1:1 to 1:1000. Competition dynamics were assessed by determining CFU/mL on selective agar plates to evaluate competition dynamics.

#### f) Screening against human-associated pathogenic bacteria

To evaluate the interaction of the reference strain of *K. oxytoca* and various known intestinal pathogenic bacteria, additional strains (**Supplementary Table 1**) were evaluated in the standard 10:1 ratio of *K. oxytoca*-to-competitor. For coculture preparation, all bacterial strains were cultured overnight in LB medium and bacterial inocula were prepared as previously described. Selection for those strains was conducted on LB agar (Carl Roth) supplemented with 10 µg/mL ciprofloxacin (Sigma-Aldrich), 50 µg/mL kanamycin (Carl Roth) or 100 µg/mL streptomycin (Sigma-Aldrich).

#### g) Screening of *A. baumannii* with human-associated commensal bacteria

To assess the interaction of the reference strain of *A. baumannii* with human-derived intestinal bacteria, various commensal bacteria were selected. A first set included strains from the KoSC, specifically *K. oxytoca*, *K. grimontii* and *K. michiganensis*. A second panel consisted of various intestinal commensal bacterial, all isolated from human feces (**Supplementary Table 1**). All strains were evaluated in the standard 10:1 ratio of commensal-bacteria-to-*A. baumannii*. For coculture preparation, all bacterial strains were cultured overnight in LB medium (Carl Roth) for the first selection and modified Gifu Anaerobic Medium (mGAM, himedia) for the commensal bacteria set. Bacterial inocula were prepared as previously described. For KoSC, coculture assay was performed according to the standardized protocol. For the intestinal commensal set, the coculture assay was set up in mGAM with and without maltose addition. Selection for listed strains was conducted on BHI agar (Carl Roth) with 10 µg/mL ciprofloxacin (Sigma-Aldrich), 50 µg/mL kanamycin (Carl Roth) or 25 µg/mL chloramphenicol (Carl Roth).

### Testing for oxygen influence on bacterial interaction

Since *A. baumannii* is a known aerobic bacterium, the influence of oxygen on its interaction with the facultative anaerobic *K. oxytoca* was assessed under anaerobic and hypoxic conditions (0.5 % oxygen to 2 % oxygen) using multiple experimental approaches.

#### a) Comparison of environmental oxygen status

The standardized coculture assay was performed as previously described in tryptone medium with and without maltose supplementation with the two representative strains, *K. oxytoca* MK01 and *A. baumannii* FR4326. Incubation was conducted at 37 °C with shaking under three distinct conditions: (i) within an anaerobic chamber maintaining hydrogen levels >2% and without oxygen, (ii) undder oxygen-limitation using the AnaeroGen system, and (iii) under aerobic conditions (∼21 % atmospheric oxygen). Following 24 hours of incubation, all cultures were processed according to the standard coculture protocol under aerobic conditions.

#### b) Assessment of bacterial interaction under hypoxic conditions

To evaluate bacterial interactions at varying oxygen levels, coculture assays were conducted in an anaerobic chamber equipped for controlled oxygen influx (Coy). Experimental condition included oxygen concentrations of 0 %, 0.5 %, 1 % or 2 % in the gas phase, while maintaining hydrogen levels >2%. The standardized coculture assay was conducted with incubation under each of those defined conditions for 24 hours, after which all subsequent steps followed the previously described coculture protocol under aerobic conditions.

#### c) Supernatant production under hypoxic conditions

To assess the impact of excreted compounds from *K. oxytoca* on *A. baumannii* suppression under oxygen limitation, supernatants of *K. oxytoca* monocultures were collected. Monocultures were scalled proportionally to match the bacterial suspension-to-medium ratio of the standardized coculture assay and incubated in sterile 50 mL falcon tubes (Sarstedt) at 37 °C under hypoxic conditions (0 %, 0.5 %, 1 %, 2 % oxygen; hydrogen > 2%). After 24 hours, bacterial cultures were removed from the anaerobic chamber and centrifuged at 3000 rcf (Heraeus) to collect the liquid phase. The supernatant was sterile-filtered and subsequently used as the incubation medium for OD_600_-standardized bacterial suspensions of *K. oxytoca* MK01 and *A. baumannii* FR4326. Each bacterial monoculture was inoculated in the supernatant at a 1:25 ratio of bacteria-to-medium in a 96-well plate (Sarstedt) and incubated at 37 °C for 24 hours under anaerobic conditions using the AnaeroGen system. After incubation, serial dilutions in PBS were plated onto non-selective LB agar, incubated aerobically at 37 °C for 24 hours and CFU/mL were calculated.

#### d) Bacterial growth under hypoxic conditions

Overnight cultures of *K. oxytoca* MK01 and *A. baumannii* FR4326, FR3462 and the type strain DSM 30007^T^ in LB medium were standardized to an OD_600_ of 0.2. Bacterial suspensions were added to tryptone base medium with and without maltose supplementation in a 1:100 ratio in a 96-well plate (Sarstedt). The plate was transferred into an anaerobic chamber with oxygen levels at 0 %, 0.5 %, 1 % or 2 %, while maintaining hydrogen levels > 2 %. The plate then was sealed with a gas-permeable foil and placed into a microplate reader (BioTek), where bacterial growth was monitored for 72 hours at 37 °C with continuous shaking.

#### e) Bacterial resilience on surfaces under hypoxic conditions

To assess the impact of oxygen on bacterial survival on surfaces, overnight cultures in LB medium of *K. oxytoca* MK01 and *A. baumannii* FR4326, FR3462 and the type strain DSM 30007^T^ were standardized to OD_600_ 1. Serial dilutions (up to 10^7^) were prepared and plated on LB agar plates. Plates were incubated at 37 °C in a temperature cabinet in an anaerobic chamber, which was set to defined oxygen levels (0 %, 0.5 %, 1 % or 2 %). Every 24 hours, a respective set of plates was removed from the chamber and, if no colonies were yet visible, incubated aerobically for an additional 24 hours at 37 °C. Colony formation was recorded and CFU/mL were calculated.

### Kinetic observation of coculture development

To assess the time line of bacterial interaction, in parallel, standardized coculture assays were started and stopped every two hours for incubation times up to 16 hours in separate boxes utilizing the AnaeroGen system. A final data set was obtained at 24 hours incubation. For every sampling timepoint, CFU/mL of bacterial competitors were determined on selective agar plates, as well as pH levels of cultures and media utilizing pH indicator strips (Merck). Additionally, at everly timepoint, from three pooled technical replicates for every biological replicate, culture liquid was pipetted into a sterile Eppendorf tube. The tubes then were spun down for five minutes at room temperature at 6000 rcf (Eppendorf) and supernatant was transferred into new sterile Eppendorf tubes. Supernatant samples then were frozen at -20 °C until collectively processed for lactic acid detection.

### Impact of acidity on bacterial survival

To assess the impact of different pH environments on bacterial survival on surfaces, overnight cultures in LB medium of *K. oxytoca* MK01 and *A. baumannii* FR4326 were standardized to an OD_600_ of one. Tryptone base media with and without maltose supplementation were prepared with installed pH from 3.5 to 9.5 with a pH meter (Mettler Toledo) using hydrocholoric acid (HCl) and sodium hydroxide (NaOH). Bacterial suspensions were added to the different media in a 1:25 bacteria-to-medium ratio in a 96-well plate (Sarstedt) for monocultural growth. The plate was incubated anaerobically in an AnaeroGen system for 24 hours at 37 °C with shaking. After incubation, serial dilutions (up to 10^7^) in PBS were prepared and plated on non-selective LB agar (Carl Roth) plates. Plates were incubated at 37 °C aerobically. Colony formation was recorded and CFU/mL was calculated.

### Impact of serine on *A. baumannii* growth

To assess the previously reported growth-supporting effect of L-serine on *A. baumannii*, overnight cultures of FR4325 in LB medium were standardized to an OD_600_ of 0.2. Tryptone base medium was supplemented with L-serine (Carl Roth) at final concentrations of 19 mM or 38 mM. To simulate an acidic environment, the pH was adjusted to 5 or 6 for test media prior to bacterial inoculation. Bacterial suspensions were added to various media each at a 1:25 ratio of bacteria-to-medium in a 96-well plate (Sarstedt) and incubated at shaking at 37 °C for 24 hours under anaerobic conditions using the AnaeroGen system with shaking. Following incubation, serial dilutions in PBS were plated onto non-selective LB agar (Carl Roth) and incubated aerobically at 37 °C for 24 hours. After plate incubation, bacterial colonies were counted and CFU/mL calculated.

### Exclusion of cell-cell contact in bacterial competition

To prevent direct physical interaction between *K. oxytoca* MK01 and *A. baumannii* FR4326, the Cerillo Coculture Duet System was employed. Bacterial cultures were incubated in tryptone base medium with and without maltose supplementation. A 0.2 µm membrane separated two wells, thereby creating two separate bacterial incubation environments while allowing the exchange of membrane permeable compounds in their shared medium. This setup ensured that cell-cell contact between bacterial cultures was inhibited. The coculture assay was performed under standardized conditions, with bacterial input parameters (OD_600_ and cell concentration) adjusted according to manufacturer’s recommendations for a well volume of 400 µL. Monocultures and cocultures were incubated at 37 °C with continuous shaking for 24 hours in an anaerobic chamber in a microplate reader (BioTek). To determine bacterial viability and potential cross-contamination, an aliquot of the input culture was diluted in phosphate-buffered saline (PBS) and plated on selective agar for colony-forming unit (CFU/mL) estimation. Following incubation, cultures were diluted in PBS under aerobic conditions and plated in dilutions on selective agar plates *A. baumannii* on LB agar + antibiotic; *K. oxytoca* on CHROMagar orientation). Plates were incubated aerobically at 37 °C overnight and bacterial colonies enumerated for calculations of CFU/mL.

### Metabolomics

*Klebsiella oxytoca* MK01 was cultured in tryptone-based medium supplemented with 10 g/L maltose in 96-well plates, following the dilutions used in the standardized coculture assay. Incubation was performed at 37 °C with shaking under aerobic conditions or in an anaerobic chamber for oxygen exclusion. Before and after 24 hours of incubation, the culture content from six wells were collected in a sterile Eppendorf tube. The pooled cultures were centrifuged at 7 000 × g for 10 minutes at 4 °C. Subsequently, 1 mL of the supernatant was transferred to a cryogenic Eppendorf tube and stored at −20 °C until further metabolomic analysis.

For extracellular metabolites, cell-free culture supernatants were diluted 1:10 with water. 50 µL of the diluted samples were mixed with 10 μL of methanol/^13^C-ribitol solution (0.8 mg/L (w/v)) and then dried in a vacuum concentrator. Derivatization by methoxyamine and N-methyl-N-(trimethylsilyl)trifluoroacetamide and subsequent GC-MS measurement (splitless and split 1:10) of metabolites were performed as described in a previous study^104^. Processing of raw data was performed as described earlier using the MetaboliteDetector software. The peak area data were normalized to the internal standard ^13^C-ribitol. ^104–106^

### Enzymatic lactic acid determination

To quantify lactic acid in supernatant samples collected at different incubation time points during the kinetic observation experiment, a commercial assay kit (K-DLATE, Megazyme) was used to separately determine D- and L-lactic acid concentrations in liquid samples. The quantification was based on two enzymatically catalysed reactions involving D-lactate dehydrogenase (D-LDH) or L-lactate dehydrogenase (L-LDH) and D-glutamate-pyruvate transaminase (D-GPT). The assay was conducted according to manufacturer’s instructions with all volumes scaled down tenfold to accommodate a 96well plate format (Sarstedt) for high-throughput analysis ^107,108^. Absorbance measurements were performed using a microplate reader (BioTek).

### *Ex vivo* coculture assay

To evaluate bacterial competition in a biologically relevant environment, the standard coculture assay was performed using previously frozen cecum content collected from mice. The cecum content was diluted 1:20 in phosphate-buffered saline (PBS) and either sterile-filtered or used unfiltered, with and without maltose supplementation from a sterile maltose stock solution. The dilution reduced animal-derived cecum content for ethical reasons, eased pipetting, and minimized physical interference in the liquid medium, allowing for more accurate bacterial culture comparisons. Cecum content from mice with various intestinal microbiota compositions was used: Germ-free (GF), specific pathogen-free (SPF) and OMM-12 with a specific microbiome, composed of twelve defined murine bacterial strains^74^. Prior to each assay, the cecum content microbiome background was tested to ensure, that it did not interfere with the enumeration of *K. oxytoca* and *A. baumannii* on respective selective agar plates. Subsequent steps in the assay followed the procedure of the standardized coculture assay.

### In vivo Galleria melonella *model*

Final instar *Galleria mellonella* larvae were used for all experiments, in accordance with general research practices for *Galleria* infection models ^53,83,109,110^. Larvae were maintained under dark conditions at room temperature and used within one week of delivery. For all setups, groups of 10 larvae were maintained in parallel to allow standardized scoring of survival and health. Scoring followed adapted methods ^76,83^ with assessments at 6, 18, 24, 48, 72, and 96 h post-infection. Non-moving and melanized larvae were removed at the respective time points.

Larvae were force-fed with 10 µL bacterial inocula prepared in PBS (Gibco) (with or without sugar supplementation) were administered orally using a PB600 dispenser (Hamilton) fitted with a blunt-end 27G needle (B.Braun). Larvae were gently stimulated until they opened their mandibles and then presented with a droplet for ingestion ^81,82^.

Across all experiments, the same bacterial strains were used*: A. baumannii* FR4326 and the tilimycin-negative probiotic candidate *K. oxytoca* MK01 *ΔnpsA*. Both strains were grown in LB medium for 4–5 h under aerobic conditions, harvested by low-speed centrifugation, washed once with PBS, and resuspended in PBS. Optical density (OD_600_) was measured, and suspensions were adjusted to OD_600_= 4. Depending on the experimental setup, further dilutions were prepared in PBS with or without maltose (20 g/L) to a final OD_600_= 2, corresponding to approximately 10^7^ CFU per feeding. This procedure ensured comparability across all experiments and standardized sugar concentrations across groups.

Two experimental designs were implemented:

a) Colonization assay. To assess bacterial colonization ability, larvae were divided into non-ampicillin-treated and ampicillin-pretreated groups. Pretreatment consisted of oral force-feeding with 10 µL of ampicillin solution (1 g/L) 24 h prior to bacterial administration, in order to reduce native microbiota and create a permissive niche. For each condition, groups comprised 10 larvae used for phenotypic scoring and 5 additional larvae harvested at 48 h post bacterial administration for quantitative analysis. Within each treatment category, larvae were assigned to the following groups: non-fed control, PBS-fed control, *A. baumannii*-fed, or *K. oxytoca*-fed. For determination of the location of bacterial colonization, resection of the gastrointestinal tract out of a subset of larvae after snap freezing and thawing was performed, and larvae body and gastrointestinal tract were seperately homogenized and plated.
b) Force-feeding with and without maltose. To evaluate sugar-dependent bacterial interactions, larvae were force-fed with bacterial suspensions prepared in PBS with or without maltose supplementation. At least 27 larvae per group were analyzed, derived from two independent larval batches. Observations were based on 2 × 10 larvae per group. Experimental procedures followed the same inoculum preparation and force-feeding method as described above, including polybacterial infections in which each bacterial strain was administered in a separate 10 µL droplet.

For quantitative recovery of bacterial burden, larvae were harvested and immediately snap-frozen in liquid nitrogen. Each larva was homogenized individually in 3 mL sterile PBS using a mechanical tissue grinder in a 15 mL screw-cap tube. Homogenization was performed on ice, and repeated if needed. The homogenizer was cleaned with ethanol and PBS between samples. Homogenates were plated on selective agar media (*K. oxytoca* on CHROMagar orientation; *A. baumannii* on LB agar with kanamycin), and CFU/larva were enumerated.

### Isolation of commensal bacteria

Stool samples from healthy human donors, participating in the LöwenKIDS cohort, were homogenized in 1 mL PBS. Serial dilutions in PBS of the homogenisate were plated on CHROMagar orientation plates. After 24 hours and 48 hours incubation at 37 °C, single colonies were picked according to differences in phenotypic appearance and streaked onto fresh CHROMagar orientation plates to generate single strain isolates. Plates were incubated again for 24 hours to 48 hours at 37 °C. Single colony streaking was repeated until pure single strain isolates were obtained and could be catalogued.

### Whole genome sequencing

To assess the taxonomy of all isolated commensal isolates and *A. baumannii* FR4326, along with FR3462, bacteria were sent for whole genome sequencing. First, genomic DNA was extracted from pelleted liquid cultures using the ZymoBIOMICS 96 MagBead DNA-Kit according to the manufacturer’s instructions. Afterward, libraries of each isolate were prepared using the Illumina DNA PCR-Free Prep and quantified with the KAPA Library Quantification Kit Illumina Platforms. Samples were pooled and sent for whole genome sequencing performed by the group of genome analysis at HZI using NovaSeq 6000 S4 Reagent Kit v1.5 (300 cycles) and targeting depth of 1 million reads per sample.

### Visualization, mathematical calculations and statistical analysis

Biolog data visualized using R studio (2024.12.1 Build 563) with packages ggplot2(3.5.1), reshape2(1.4.4), circlize(0.4.16)^111^ & ComplexHeatmap(2.22.0), tidyverse (2.0.0), RColorBrewer(1.1-3), dendextend(1.19.0), dplyr(1.1.4).

Competitive indices (CI) were calculated according to reported models of mixed bacterial infections ^112,113^. CI is defined as the ratio of the products of recovered CFU/mL for each bacterium and their respective competitor’s input.

Experimental results were analyzed for statistical significance using GraphPad Prism (v10.4) (GraphPad Software Inc.). *P*-values were calculated by non-parametric Mann-Whitney U test or Kruskal-Wallis test comparison of totals between groups. *P*-values lower than 0.05 were considered significant: \**p* <0.05, \*\**p*<0.01, \*\*\**p*<0.001 \*\*\*\**p*<0.0001. Further description of each statistical test is supplied in the figure legends.

## Data availability

All data generated or analyzed during this study are included in the published article or its supplementary information. Explanatory graphics were created in BioRender. Supplementary Data are provided with this publication. Additional data are available from the corresponding author upon reasonable request.

## Supplementary Data

Supplementary Table 1: Bacterial strains used in this study

Supplementary Table 2: GC-MS peak intensity data for MK01 culture analysis in tryptone base medium with maltose supplementation

Supplementary Table 3: Adapted Scoring Table for *Galleria* model

Supplementary Figure 1: Influence of cocultivation with *K. oxytoca* on the growth of *A. baumannii* and related *Acinetobacter* spp. across diverse media and conditions

Supplementary Figure 2: Detailed information of oxygen- and maltose-dependent growth dynamics of *A. baumannii* and *K. oxytoca* in mono- and coculture

Supplementary Figure 3: Growth dynamics of *K. oxytoca* during suppression of *A. baumannii*

Supplementary Figure 4: Influence of pH, metabolites and lactata isomers on *K. oxytoca-A. baumannii* interactions under oxygen-limited conditions

Supplementary Figure 5: Growth dynamics of *K. oxytoca* and commensal bacteria during interaction with *A. baumannii*

## Acknowledgements

We sincerely thank Agata Bielecka, Achim Gronow and Karin Paduch for their technical support in experiments and maintaining the lab infrastructure. Our gratitude also goes to Gesa Martens for ensuring smooth operations at our cooperator’s facility. We appreciate the valuable assistance of Anastasia Giese and Dorothee-Sophie Börstling.

## Author contributions

K.A.W. and T.S. conceived and designed the experiments. K.A.W. and M.W. conducted the experiments and analyzed the data. Data analyses was conducted by K.A.W., except for whole genome sequencing analysis, conducted by T.R.L.. M.N.S. provided metabolomic analysis and biochemical insights. M.W. contributed to not included data in the conceptual framework. L.O., along with V.A.J.K., provided reference strains for this study. E.A. and L.E. supplied genetically modified bacterial strains. T.S., L.O., J.O. and D.S. provided critical support and expert advice. Additional bacterial strains used in this study were obtained from M.H., V.A.J.K., L.K., N.P., S.G., B.A., J.O., D.S.. All authors reviewed and approved the final manuscript.

## Contact for reagent and resource sharing

Further information and requests for resources and reagents should be directed to and will be fulfilled by the Lead Contact, Prof. Dr. Till Strowig

## Ethics declaration

We support inclusive, diverse and equitable conduct of research.

