## Supplementary Figures for "Metabolites produced by gut bacteria under anoxic conditions drive the suppression of Acinetobacter baumannii"

**Supplementary Fig. 1**

**a** *A. baumannii* FR4326 was cultured alone or in coculture with *K. oxytoca* MK01 (10:1 ratio) in various media ± 10 g/L maltose. Respective media are tryptone base medium (TBM), Minimal medium 9 (MM9), Luria-Bertani medium (LB) and modified Gifu Anaerobic Medium (mGAM). The growth of *A. baumannii* was determined by selective plating. Data are mean CFU/ml ± SD for n= 5 biological replicates.

**b, c** Coculture of varying *Klebsiella*: *Acinetobacter* ratios of *A. baumannii* FR4326 and *K. oxytoca* MK01 grown in mono- and coculture in TBM + 10 g/L maltose. The growth of *A. baumannii* (**b**) and *K. oxytoca* (**c**) was determined by selective plating. Data are means ± SD for n= 3 biological replicates.

**d** *A. baumannii* FR4326 was grown alone or in coculture with *K. oxytoca* MK01 in TBM supplemented with increasing maltose concentrations. The growth of *A. baumannii* was determined by selective plating. Data is shown as mean ± SD for n= 4 biological replicates.

**e - k** *K. oxytoca* MK01 was cocultured with the indicated *Acinetobacter* spp. strains in TBM ± 10 g/L maltose. (**e**) *A. baumannii* strains from the gastrointestinal tract (GIT), respiratory tract (RT), and skin wounds (W) (**f - k**) GIT isolates of *A. baumannii* (Ab), *A. nosocomialis* (An), and *A. pittii* (Ap). The growth of *Acinetobacter* strains and *K. oxytoca* was assessed by selective plating (*Acinetobacter* spp. on streptomycin **f**, on kanamycin **h**, ciprofloxacin **j**; *K. oxytoca* on Chrom agar orientation **g**, **i**, **k**). Data are the mean ± SD for n= 3 (**e**) and n= 4 (**f - k**) biological replicates.

All coculture assays were incubated under oxygen-limited conditions using the AnaeroGen system. The detection limit (dotted line) and the inoculum (mixed dashed line) are indicated. Statistical significance was determined using Mann-Whitney U test (a, d, f – k) (*p < 0.05, **p<0.01) and Kruskal-Wallis test (**b**) with *p= 0.00198 for 10:1 and 1:1 ratios(*p < 0.05).

**Supplementary Figure 1**

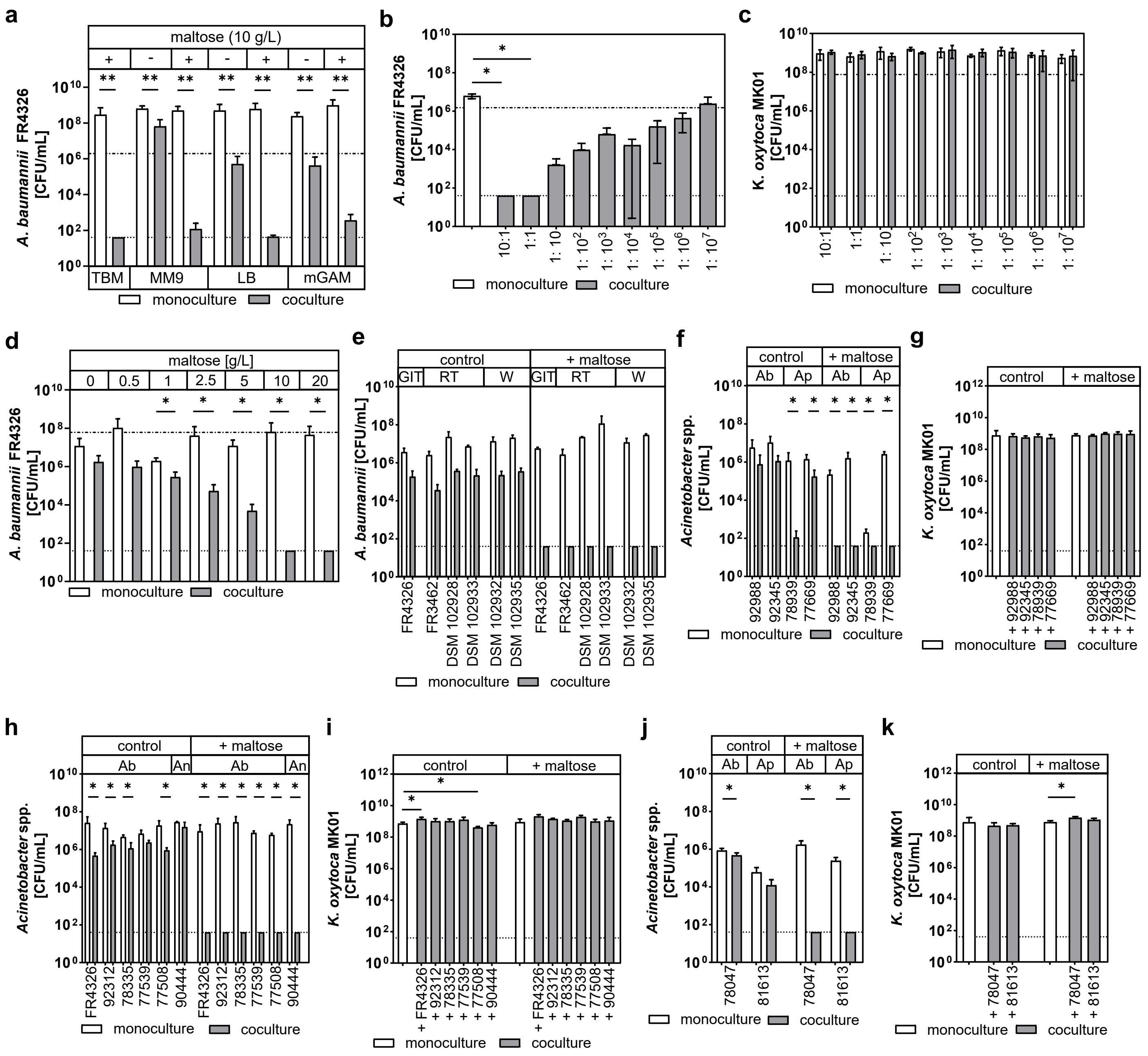

**Supplementary Fig. 2**

**a** *A. baumannii* FR4326 was cultured alone or in coculture with *K. oxytoca* MK01 in TBM ± 10 g/L maltose under different oxygen conditions: anaerobic chamber (0 % oxygen), AnaeroGen system, or aerobic conditions. The growth of *K. oxytoca* MK01 was assessed through selective plating. Data are shown as mean ± SD for n= 3 biological replicates.

**b** *A. baumannii* FR4326 was cultured alone or in coculture with *K. oxytoca* MK01 in TBM ± 10 g/L maltose under conditions with defined oxygen concentrations (0 % to 2 % oxygen). The growth of *K. oxytoca* MK01 was assessed through selective plating. Data are shown as mean ± SD for n= 4 biological replicates.

**c – j** Growth curves, based on optical density $OD_{600}$ measurements over 72 hours for *K. oxytoca* MK01, *A. baumannii* DSM 30007$^T$, FR3462 and FR4326 grown in TBM ± maltose under hypoxic conditions (0 % to 2 % oxygen) for n= 4 biological replicates.

**k – n** Recovery of $OD_{600}$ 1 cultures of *K. oxytoca* MK01, *A. baumannii* DSM 30007$^T$, FR3462 and FR4326, plated on LB agar and incubated for 0 to 120 hours under hypoxic conditions (0 % to 2 % oxygen). Data are shown as mean ± SD for n= 4 biological replicates. Colonies were counted after hypoxic incubation directly for all bacterial strains (*) or after additional 24 hours aerobic incubation to increase colony sizes to countable dimensions. Additional incubation information can be found in the methods.

The detection limit (dotted line) and the inoculum (mixed dashed line) are indicated. Statistical significance was determined using non-parametric Kruskal-Wallis test (**a**) (*p= 0.0488) and Mann-Whitney U test (**b**) (*p < 0.05).

### Supplementary Figure 2

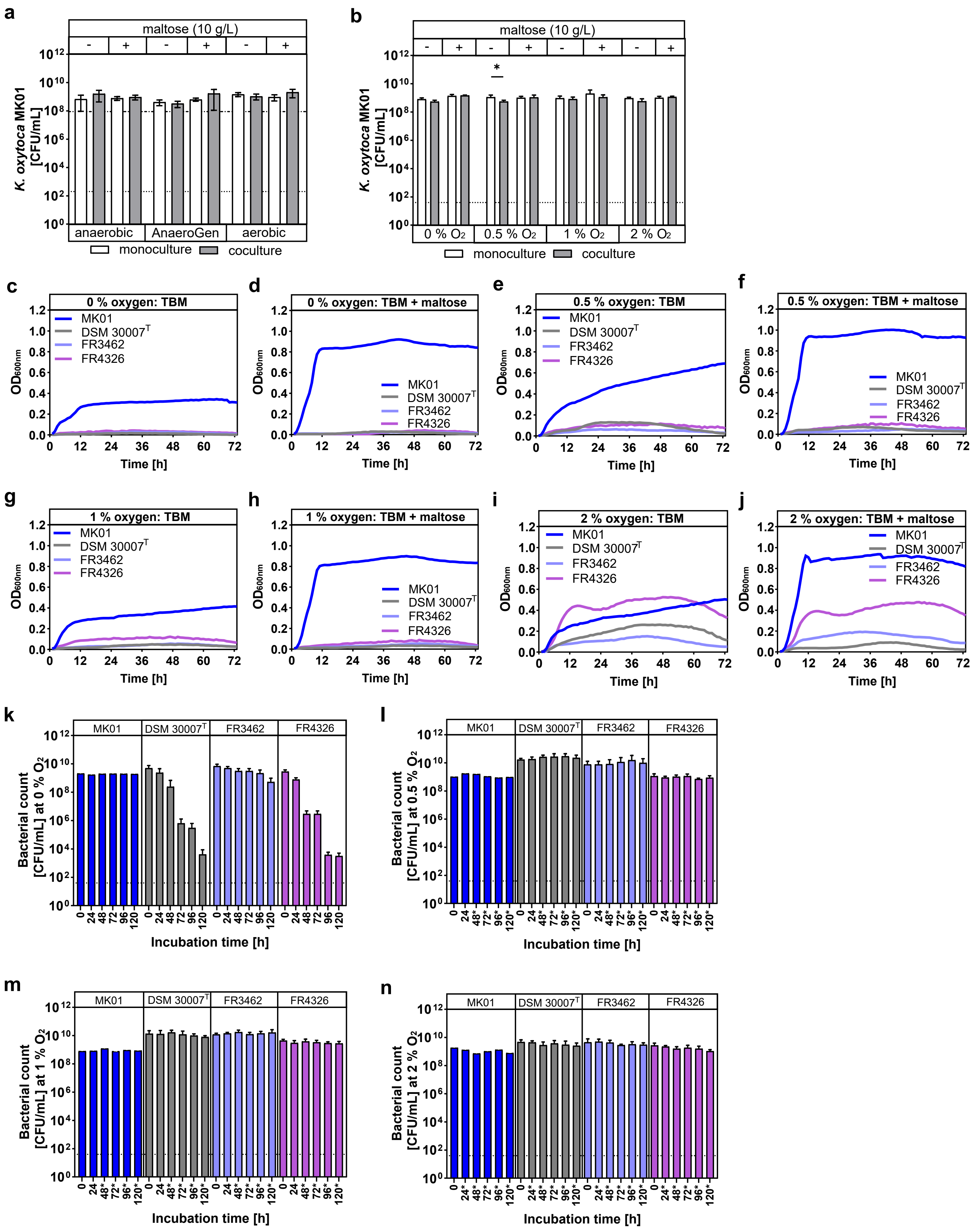

**Supplementary Fig. 3**

**a** *A. baumannii* FR4326 was grown alone or in coculture with *K. oxytoca* MK01 wt, Δ*lacZ,* and Δ*sacX* in TBM ± 10 g/L maltose, sucrose or lactose. The growth of *K. oxytoca* was assessed by selective plating. Data are shown as mean ± SD for n= 4 biological replicates.

**b** Longitudinal analysis of growth of *K. oxytoca* MK01 and *A. baumannii* FR4326 grown alone and in coculture in TBM + 10 g/L maltose. Parallel assays were stopped every 2 hours between 0 and 16 hours of incubation to collect supernatant for lactate quantification, growth enumeration (**b**), and pH measurement. The growth of *K. oxytoca* (**b**) was assessed by selective plating. Data are presented as mean ± SD for n= 3 biological replicates.

**c** *K. oxytoca* MK01 was cultured in TBM with and without maltose for 24 hours under hypoxic conditions (0 % to 2 % oxygen) in n= 4 biological replicates with incubation media controls. Steril-filtered spent media and controls, as well as fresh media were then used to culture initial $OD_{600}$ 1 *K. oxytoca* MK01 under AnaeroGen conditions and plated on LB agar plates to display sole influence of supernatant on standardized cultures. Data are shown as mean ± SD for n= 4 biological replicates

All coculture assays were incubated under oxygen-limited conditions using the AnaeroGen system if not mentioned otherwise. The detection limit (dotted line) is indicated. Statistical significance was determined using Mann-Whitney U test (**a, c**) and Wilcoxon matched-pairs signed rank test (**b,** ns) (*$p < 0.05$).

**Supplementary Figure 3**

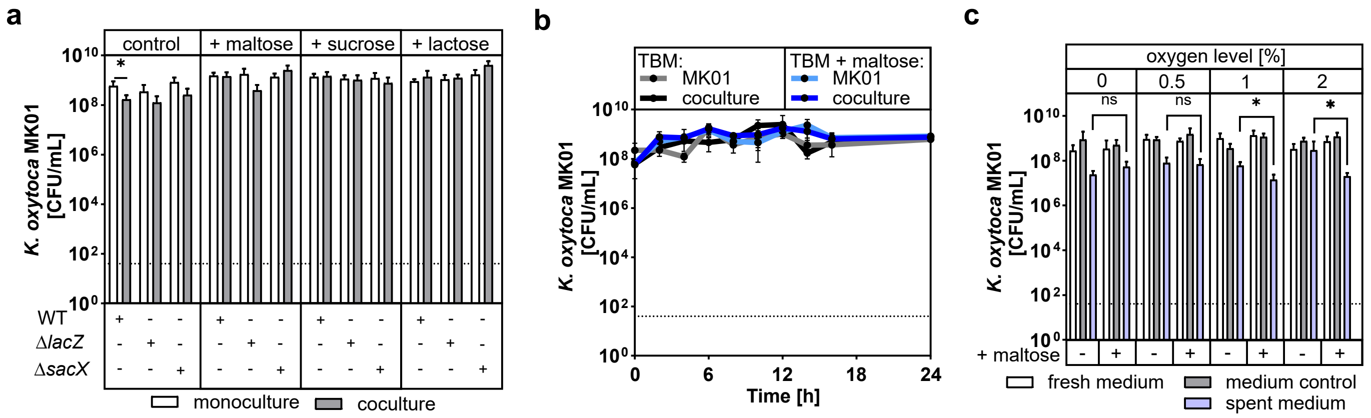

**Supplementary Fig. 4**

**a - d** *A. baumannii* FR4326 was grown alone or in coculture with *K. oxytoca* MK01 in TBM ± 10 g/L maltose and Good's buffer supplementation for pH stabilization. Added buffer in concentrations of 50mM to 200 mM were: 2-morpholin-4-ylethanesulfonic acid (MES) at pH 5, 3-(morpholin-4-yl)propane-1-sulfonic acid (MOPS) at pH 7, 3- {[1,3- Dihydroxy- 2- (hydroxymethyl)propan-2-yl]amino}propane-1-sulfonic acid (TAPS) at pH 9. The growth of *A. baumannii* and *K. oxytoca* was assessed by selective plating. Data are shown as mean ± SD for n= 2 biological replicates.

**e** GC-MS metabolomic analysis of *K. oxytoca* MK01 culture supernatant in TBM + 10 g/L maltose. Sample were collected before incubation (BI), and after 24 hours of incubation under oxygen-rich (ox) or anoxic (anox) conditions in an anaerobic chamber. Relative abundances of detected compounds are shown.

**f – h** *A. baumannii* FR4326 was grown alone or in coculture with *K. oxytoca* MK01 in TBM with the addition of increasing concentrations 0 to 98 mM cadaverine (**f**), 0 to 56 mM glucono- 1,5- lactone (**g**) or 0 to 77 mM pyroglutamate (**h**). The growth of *A. baumannii* was assessed by selective plating. Data are shown as mean ± SD for n= 3 biological replicates.

**i, j** *A. baumannii* FR4326 grown alone and in coculture was assessed for response to serine exposition. (**i**) $OD_{600}$ 0.2 standardized monocultures of *A. baumannii* were grown in TBM with 0 to 38 mM serine addition with initial media pH set from pH 5 to 7. The growth of *A. baumannii* was assessed by plating on LB agar plates. (**j**) *A. baumannii* FR4326 was grown alone or in coculture with *K. oxytoca* MK01 in TBM + 10 g/L maltose and 38 mM serine with initial media pH of 5 or 7.5. The growth of *A. baumannii* was assessed by selective plating. Data (**i**, **j**) are shown as mean ± SD for n= 3 biological replicates.

**k** Compositional graph displaying *A. baumannii* FR4326 grown in mono- and coculture with *K. oxytoca* MK01 in TBM + 10 g/L maltose supplementation and the addition of either 5.6 mM glucono-1,5-lactone, 38 mM serine, 9.8 mM cadaverine or 7.7 mM pyroglutamate. *A. baumannii* was assessed by selective plating. Data are shown as mean ± SD for n= 3 biological replicates.

**l, m, n** Accumulation of lactic acid in culture supernatant over time was measured by enzymatic detection. Supernatant was collected during longitudinal growth analysis of *K. oxytoca* MK01 and *A. baumannii* mono- and cocultures in n = 3 biological replicates. Data is shown as mean ± SD.

**o - r** *A. baumannii* FR4326 was grown alone or in coculture with *K. oxytoca* MK01 in TBM ± 10 g/L maltose and the addition of D-, D/L- or L-lactate at concentrations of 11mM or 111 mM. The growth of *A. baumannii* and *K. oxytoca* was assessed by selective plating. Data are shown as mean ± SD for n= 3 biological replicates.

All mono- and coculture assays were incubated under oxygen-limited conditions using the AnaeroGen system if not mentioned otherwise. The detection limit (dotted line) and the inoculum (mixed dashed line) are indicated.

### Supplementary Figure 4

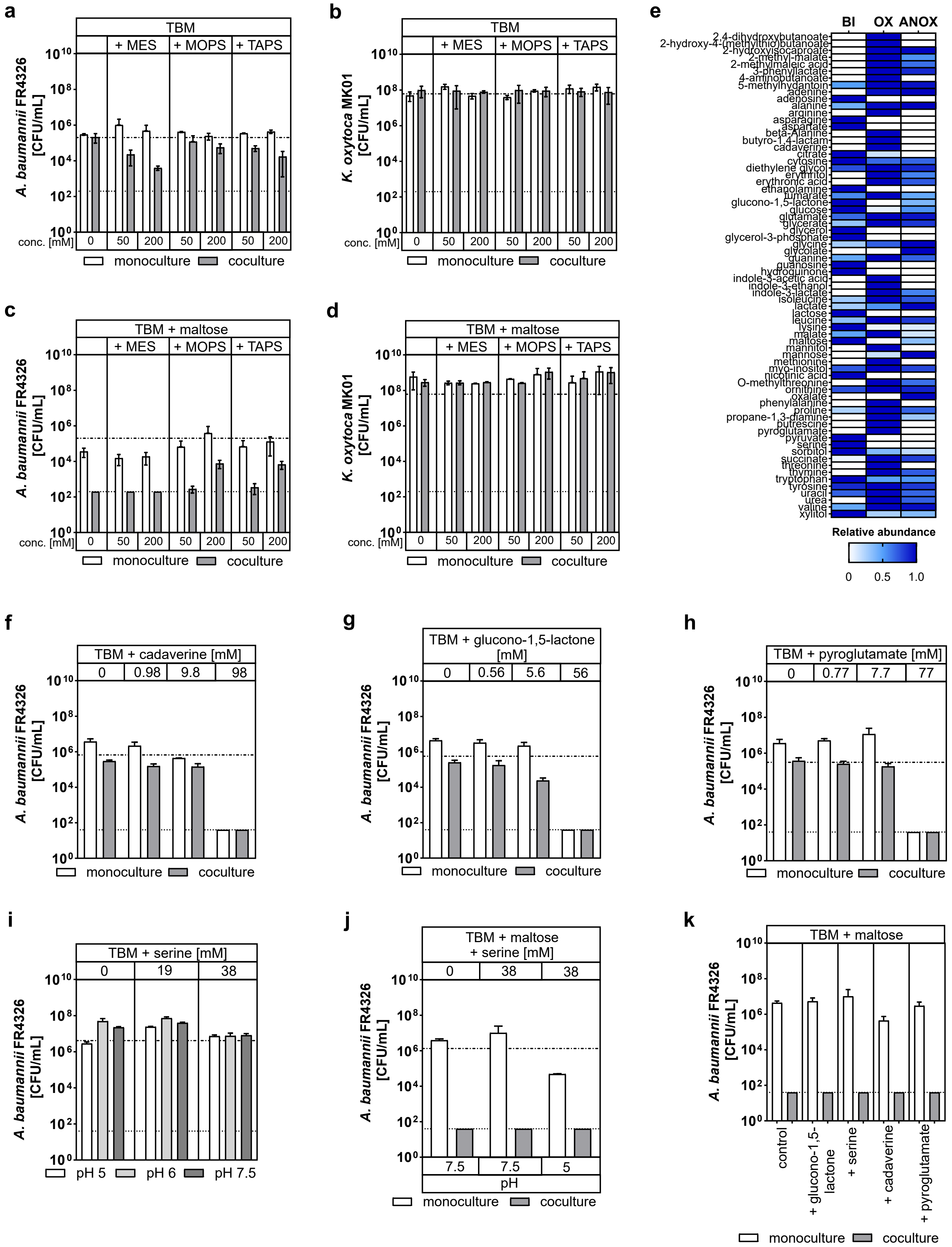

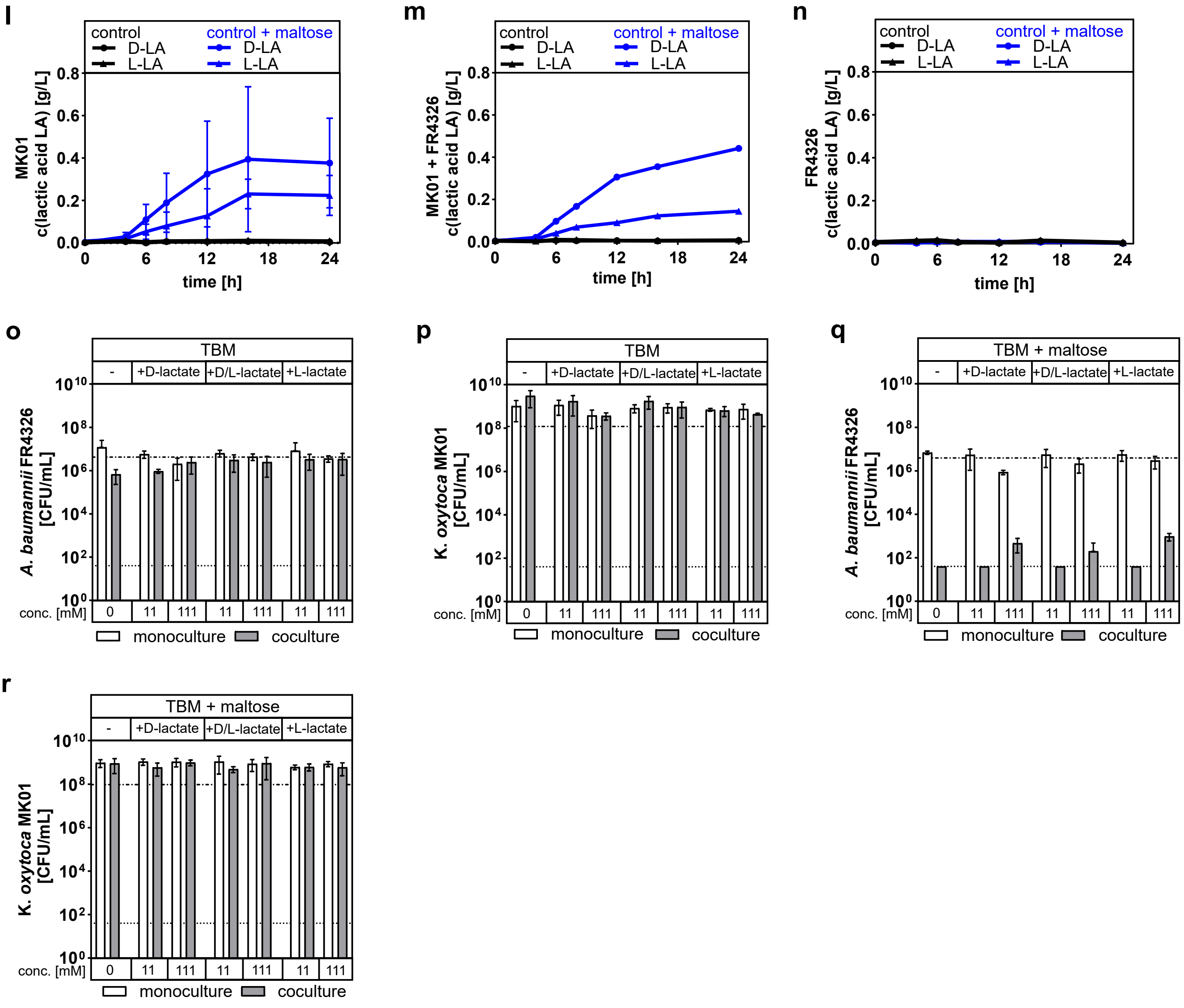
l
control
D-LA
L-LA
control + maltose
D-LA
L-LA
MK01
c(lactic acid LA) [g/L]
0.0
0.2
0.4
0.6
0.8
0
6
12
18
24
time [h]
m
control
D-LA
L-LA
control + maltose
D-LA
L-LA
MK01 + FR4326
c(lactic acid LA) [g/L]
time [h]
n
control
D-LA
L-LA
control + maltose
D-LA
L-LA
FR4326
c(lactic acid LA) [g/L]
time [h]
o
TBM
-
+D-lactate
+D/L-lactate
+L-lactate
A. baumannii FR4326
[CFU/mL]
conc. [mM]
0
11
111
11
111
11
111
monoculture
coculture
p
TBM
K. oxytoca MK01
[CFU/mL]
conc. [mM]
monoculture
coculture
q
TBM + maltose
A. baumannii FR4326
[CFU/mL]
conc. [mM]
monoculture
coculture
r
TBM + maltose
K. oxytoca MK01
[CFU/mL]
conc. [mM]
monoculture
coculture

**Supplementary Fig. 5**

**a** *A. baumannii* FR4326 was cultured alone or in coculture with various intestinal commensal species in mGAM ± 10 g/L maltose. The growth of commensal bacteria was assessed through selective plating. The mean ± SD for n= 3 biological replicates is displayed. Commensal strains are: *K. oxytoca* MK01 (Ko), *Lactobacillus paracasei* LK 355 Iso 2 (Lp), *Citrobacter braakii* LK 280 Iso 12 (Cb), *Enterobacter ludwigii* LK 290 Iso 14 (El), *Enterococcus faecium* LK 290 Iso 7 (Ef), *Lactococcus lactis* LK 370 Iso 1 (Ll), *Streptococcus oralis* LK 323 Iso 5 (So).

**b, c** *A. baumannii* FR4326 was grown alone or in coculture with *K. oxytoca* MK01 in TBM, as well as murine cecum content with and without 10 g/L maltose. Germ-free (GF), specific-pathogen-free (SPF) or OMM-12 – carrying cecum content was used either filtered (**b**) or unfiltered (**c**) in a 1:20 dilution in PBS. The growth of *K. oxytoca* was assessed through selective plating. Data is displayed as the mean ± SD for n= 6 biological replicates of bacteria, cultured in n= 2 biological replicates of cecum content.

**d, e, f** Establishment of the *Galleria mellonella* intestinal colonization model. Larvae were orally force-fed with *A. baumannii* (*Ab*), tilimycin-negative strain *K. oxytoca* MK01 Δ*npsA* (*Ko*), PBS, or left unfed (control), with or without prior ampicillin pretreatment. Survival was monitored over 120 h (**d**), and a fitness index integrating survival, activity, melanization, and cocoon/pupa formation was calculated (**e**). Bacterial recovery 48 h post bacterial force-feeding was assessed by CFU enumeration per larva, with snap-freezing and resection testing ensuring that gastrointestinal bacterial colonization was measured (**f**) (whole larval body WB, body without gastrointestinal tract WB-GI, only larval gastrointestinal tract GI).

**g, h** *Galleria mellonella* larvae were force-fed with either *A. baumannii* FR4326 (*Ab*) or *K. oxytoca* (*Ko*) alone or in combination (*Ab* + *Ko*), with or without maltose supplementation. Bacterial recovery from larval homogenates was determined 48 h post-infection by selective plating. Data represent mean ± SD from at least 27 larvae per condition (two independent bacterial culture replicates). Pairwise statistical differences were analyzed using the Mann–Whitney U test (****p < 0.0001). Overall statistical significance was determined by Kruskal–Wallis test (****p < 0.0001). Fitness of an observation group (without sampling) of two times 10 larvae was assessed according to the described adapted Scoring Model.

All coculture assays were incubated under quasi-anaerobic conditions using the AnaeroGen system if not indicated otherwise. The detection limit (dotted line) and the inoculum (mixed dashed line) are indicated. Statistical significance was determined using Mann-Whitney U test (*p < 0.05) with no statistical significance was found for (**b**) and (**c**).

### Supplementary Figure 5

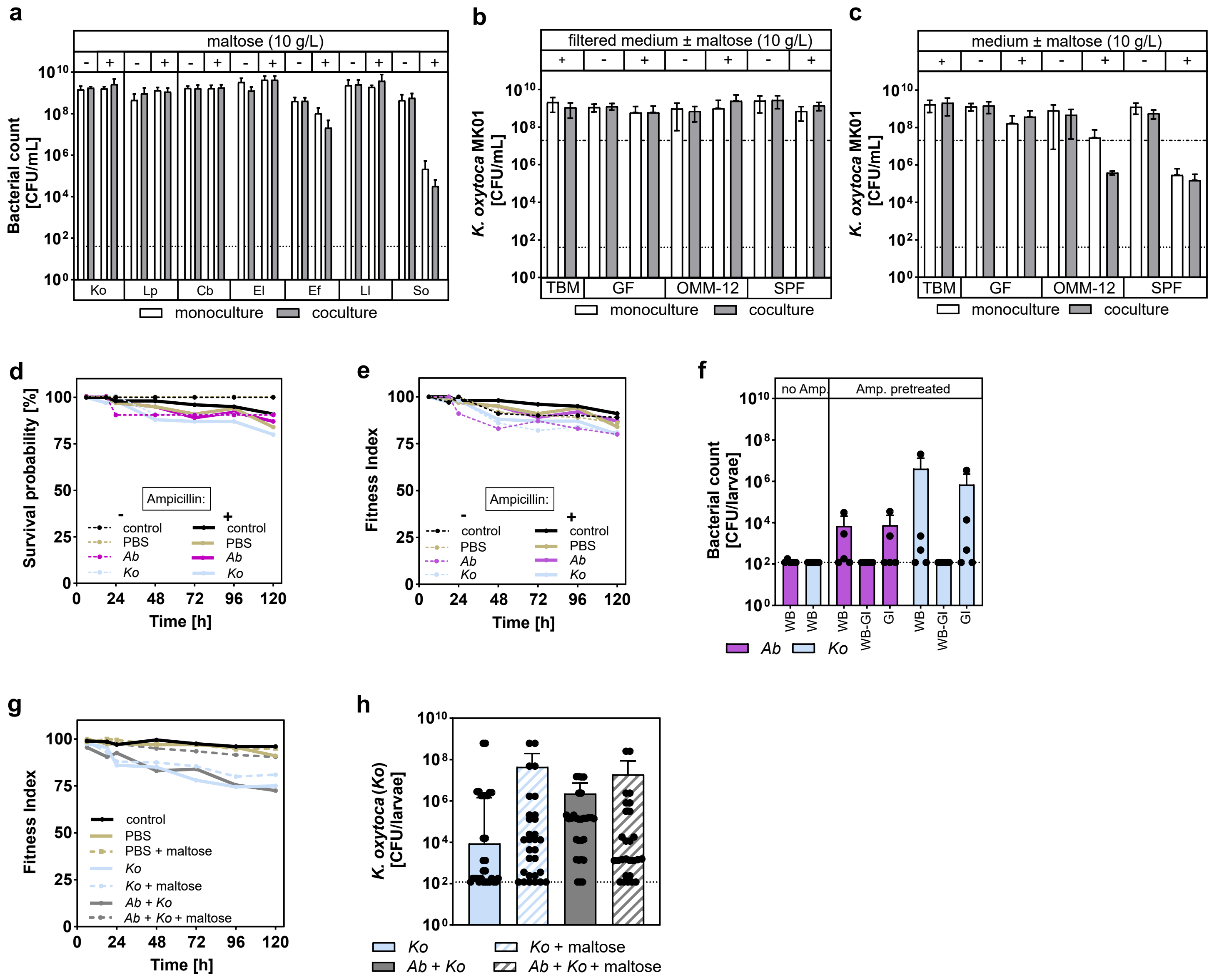
